# Self-assembly and structure of a clathrin-independent AP-1:Arf1 tubular membrane coat

**DOI:** 10.1101/2022.05.23.493093

**Authors:** Richard M. Hooy, Yuichiro Iwamoto, Daniel Tudorica, Xuefeng Ren, James H. Hurley

**Affiliations:** Department of Molecular and Cell Biology, University of California Berkeley; Berkeley CA 94720, USA; California Institute for Quantitative Biosciences, University of California, Berkeley, CA, 94720, USA; Graduate Group in Biophysics, University of California, Berkeley, CA, 94720, USA; Helen Wills Neuroscience Institute, University of California, Berkeley, Berkeley, CA 94720, USA

## Abstract

The AP adaptor complexes are best known for forming the inner layer of clathrin coats on spherical vesicles. AP complexes also have many clathrin-independent roles in tubulovesicular membrane traffic, whose structural and mechanistic basis has been a mystery. HIV-1 Nef hijacks the AP-1 complex to sequester MHC-I internally, evading immune detection. We found that AP-1:Arf1:Nef:MHC-I forms a coat on tubulated membranes in the absence of clathrin, and determined its structure by cryo-ET. The coat assembles both laterally and axially *via* an Arf1 dimer interface not seen before. Nef recruits MHC-I, but is not essential for the underlying AP-1:Arf1 lattice. Consistent with a role for AP-1:Arf1 coated tubules as intermediates in clathrin coated vesicle formation, AP-1 positive tubules are enriched in cells upon clathrin knockdown, with or without Nef. Nef localizes preferentially to AP-1 tubules in cells, explaining how Nef can sequester MHC-I. The coat contact residues are conserved across Arf isoforms and across the Arf-dependent AP adaptors AP-1, 3, and 4. These findings reveal that AP complexes can self-assemble with Arf1 into tubular coats in the absence of clathrin or other scaffolding factors. The AP-1:Arf1 coat defines the structural basis of a broader class of tubulovesicular membrane coats, as an intermediate in clathrin vesicle formation from internal membranes, and as a MHC-I sequestration mechanism in HIV-1 infection.

---

Membrane and secretory proteins and lipids are distributed to their destinations within eukaryotic cells by the trafficking of tubular and vesicular structures. Tubular and vesicular structures are organized by proteinaceous vesicle coats, which include clathrin and its AP adaptor complexes, COPI, COPII, and retromer (Dell’Angelica and Bonifacino, 2019). Clathrin and COPI are vesicular coats (Faini et al., 2013), retromer is a tubular coat (Kovtun et al., 2018), and COPII can assemble in both tubular and vesicular geometries (Faini et al., 2013). The AP adaptor complexes AP-1 to -5 function in a vast array of cellular process, of which some, but not others, depend on clathrin. The AP adaptor protein complexes AP-1 and AP-2 form the inner layer of clathrin-coated vesicles (CCV), connecting interior lipids and cargoes to the exterior clathrin cage. Despite that clathrin itself does not form tubular coats, there have been many reports of AP complexes on tubular structures, including the typically clathrin-dependent AP-1 (Braun et al., 2007; Haberg et al., 2008; Janvier and Bonifacino, 2005). AP-3 is principally localized to tubular structures and, while it can associate with clathrin, AP-3 is functionally independent of it (Peden et al., 2004; Vowels and Payne, 1998). AP-4 does not interact with clathrin at all (Dell’Angelica et al., 1999; Hirst et al., 1999). These observations suggest that AP-1, -3, and -4 must have some fundamental ability to assemble on tubular membranes. Despite extensive structural studies of the AP and related complexes (Faini et al., 2013; Jackson et al., 2012), the basis for the formation of tubular and clathrin-independent assemblies is unknown.

One example of such a coat assembly might be the AP-1-dependent sequestration of MHC-I by the HIV-1 accessory protein Nef. Downregulation of the antigen presentation complex MHC-I is a ubiquitous strategy employed by viral pathogens to subvert host CD8^+^ T cell mediated elimination of virally infected cells (van de Weijer et al., 2015). The human and simian immunodeficiency viruses, HIV-1, HIV-2 and SIV, encode an accessory factor, Nef, that achieves this critical task as one of its various functions (Buffalo et al., 2019; Collins and Collins, 2014; Dekaban and Dikeakos, 2017; Ramirez et al., 2019; Sauter and Kirchhoff, 2018; Staudt et al., 2020). Nef targets MHC-I by hijacking the clathrin adaptor complex AP-1, along with its small GTPase activator, Arf1. This reroutes mature MHC-I molecules to the perinuclear region and/or endosomal sorting network instead of the plasma membrane (Kasper et al., 2005; Le Gall et al., 1998; Lubben et al., 2007; Roeth et al., 2004; Wonderlich et al., 2011). MHC-I is eventually degraded at the lysosome in a clathrin-dependent mechanism (Roeth et al., 2004; Schaefer et al., 2008). Hijacked AP-1 and MHC-I form long-lived internal membrane-associated assemblies that sequester MHC-I away from the plasma membrane (Blagoveshchenskaya et al., 2002; Dirk et al., 2016; Janvier et al., 2003a; Le Gall et al., 1998; Tavares et al., 2020). The nature of the MHC-I-sequestering assemblies has remained obscure.

AP-1, and the four other human AP complexes (2 through 5), share a conserved heterotetrameric architecture consisting of a small (σ), medium (μ) and two large subunits; AP-1 consists of σ1, μ1, γ and β1 (Owen et al., 2004) (Fig. 1A). APs are allosterically regulated and couple conformational states to cargo binding, membrane targeting and recruitment of clathrin (Canagarajah et al., 2013; Jackson et al., 2010). In the locked conformation, AP-1 is primarily cytosolic and has a low affinity for phosphatidylinositol 4- phosphate (PI(4)P) and cargo molecules. To become activated, myristoylated Arf1 in its GTP-bound state recruits AP-1 to target membranes at the TGN and endosomes, and stabilizes the unlocked conformation of AP-1 (Ren et al., 2013). The unlocked conformation binds cargo bearing the consensus YxxΦ motif at the μ domain and the more variable dileucine cargo motifs, (D/E)xxxL(L/I/M), at the σ domain (where ‘x’ is any amino acid and Φ is any bulky hydrophobic residue). Once bound to and stabilized by cargo and myristoylated Arf1, AP-1 recruits clathrin through conserved clathrin binding motifs encoded within the extended C-terminal appendage of the β1 domain (Kelly et al., 2014). AP-1 localizes to membrane compartments enriched in PI(4)P (Wang et al., 2003), including the trans-Golgi network (TGN) and endosomal system. However, Arf1 is the major determinant of subcellular localization and activation of AP-1 (Stamnes and Rothman, 1993; Traub et al., 1993). AP-3 (Ooi et al., 1998) and the clathrin-independent AP-4 (Boehm et al., 2001) are also Arf1-dependent. Thus, AP-1, -3, and -4 comprise a subset of Arf-dependent AP complexes that are likely to be governed by common recruitment and activation principles.

**Figure 1.**
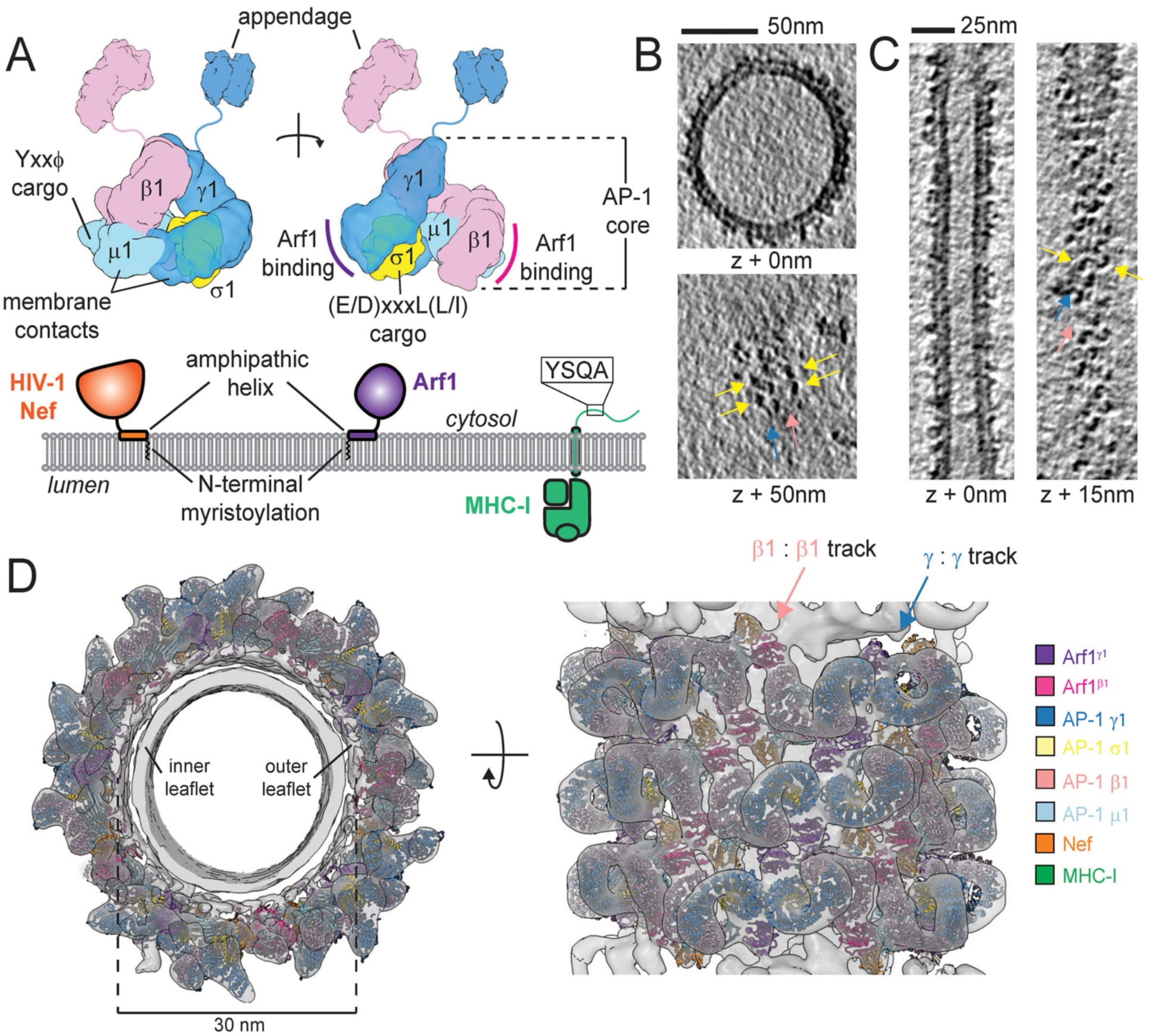
AP-1, Arf1 and Nef form an ordered coat on MHC-I^cyto^-membranes. (A) Cartoon overview of AP-1, Arf1, Nef and MHC-I. Functional sites and their associated domains are labeled. (B) Representative cryo-electron tomographic slices through an AP-1^βFL^:Arf1_myr_:Nef_myr_ coated vesicle incorporated with MHC-I^cyto^ lipopeptide. Yellow arrows indicate AP-1 density. Dark blue arrows denote tracks of AP-1 γ : γ juxtaposed molecules. Light pink arrows denote tracks of AP-1 β1 : β1 juxtaposed molecules. (C) Representative cryo-electron tomographic slices through an AP-1^βFL^:Arf1_myr_:Nef_myr_ coated tube incorporated with MHC-I^cyto^ lipopeptide. Arrows are as indicated in (B). (D) Reconstructed map and model of a continuous AP-1:Arf1:Nef coat on a section of tubular membrane. Cross-sectional view of the coat down the tube axis (left) and surface view of the coat (right). Arrows are as described in (B). Fitted AP-1, Arf1 and Nef are color coded to match the depiction in (A). Arf1 is colored pink and purple to denote association with AP-1 β1 and γ, respectively.

In the course of reconstituting the Nef-sequestered state of AP-1 and MHC-I, we noticed that AP-1:Arf1:Nef:MHC-I could form a continuous tubular protein coat on membranes in the absence of clathrin. How AP-1, -3, and -4 might organize tubular coats is a long-standing question. Viruses are often observed to “over-drive” cellular processes in the course of hijacking them, providing windows into fundamental aspects of cell biology that would be hard to obtain otherwise. Given the potential significance for tubular coat organization across eukaryotes, we determined the structure of the coat by cryo-electron tomography (cryo-ET) and subtomogram averaging (STA). The resulting structure suggests a paradigm for clathrin-independent tubular membrane coating and stabilization by the Arf1-dependent subset of AP complexes.

## Results

### AP-1, Arf1 and Nef self-assemble to form a tubular coat on membranes

The cytosolic tail of MHC-I subtypes A and B contains the consensus sequence Y(S/T)QA, which does not normally recruit AP-1 because of the presence of an Ala in place of the requisite larger hydrophobic residue of the YxxΦ motif (Noviello et al., 2008). Nef converts this non-substrate into a neo-substrate by complementing the suboptimal cargo motif (Jia et al., 2012). Nef’s tandem PxxP motif and key electrostatic interactions drive recognition of the MHC-I tail onto the canonical tyrosine motif cargo binding site of μ1 (Jia et al., 2012). To reconstitute MHC-I downregulation on membranes in vitro, liposomes containing 3 mol % MHC-I^cyto^ lipopeptide in a trans-Golgi network (TGN)-like lipid composition were generated by extrusion through a 50 nm filter. Liposomes were incubated with AP-1, myristoylated Arf1 (Arf1^myr^) and myristoylated HIV-1 NL4-3 Nef (Nef^myr^) (Fig. 1A, S1A). Arf1 nucleotide exchange was stimulated by pre-incubating Arf1^myr^, GTP and EDTA at 37^°C^ for 20 minutes followed by the addition of a molar excess Mg^2+^ over EDTA. The resulting samples were vitrified and imaged by cryo-EM.

The cryo-EM images revealed a dense protein coat on the surface of membranes (Fig. S1B). The protein coat was present on spherical vesicles, amorphous membranes, and membrane tubes, indicating that AP-1 binds to membranes of varying curvatures. To investigate the spatial distribution of AP-1 on the surface of membranes, we collected tilt series and reconstructed tomograms of the coated membranes (Fig. S2, Table S1). The tomograms show the protein coat forms a nearly contiguous layer on most membranes (Fig. 1B, C, Fig. S1C). The tomograms revealed a striking repeating density pattern consistent with a regular lattice, most obviously on tubular membranes (Fig. 1C, Fig. S1C). The patterned array was present on a range of tube diameters and was most obvious on relatively straight tube sections, and to a lesser extent on spherical vesicles (Fig 1C, Fig. S1C) and flat membrane sheets. We determined the structure of the assembly using STA. We focused on AP-1 assemblies on tubulated membranes. 72 tilt-series were collected and reconstructed into tomograms for STA (Table S1, S2). Of the 284 annotated tubes, narrow tubes were highly enriched relative to wide tubes (Fig. S1E).

Subtomograms were pooled based on tube diameter and analyzed separately to maximize structural homogeneity. Aligned and averaged subtomograms from narrow tubes (20-30 nm diameter) resulted in a 20 Å resolution map which could be fit unambiguously with atomic models of the hyper-unlocked AP-1^core^ (Jia et al., 2014; Morris et al., 2018) (Fig. 1D, Fig. S2, S3). The coat has helical symmetry with a 10 nm pitch and 10.5 AP-1 protomers per turn (Table S4). Globular densities adjacent to the fitted AP-1 heterotetramers showed density extending downward toward the membrane surface suggesting the densities belonged to the N-termini and globular domains of Arf1 and/or Nef. Placement of AP-1^core^ models (PDB: 6cm9) (Morris et al., 2018) within the lattice revealed two alternating tracks of AP-1 dimers propagating along the tube axis; one dimer pair created by γ : γ, the other by β1 : β1 contacts.

### The AP-1 coat is connected by two Arf1 bridges

The dimeric interfaces connecting the protomers laterally and axially were investigated by STA. The β1:β1- and γ:γ-juxtaposed AP-1 dimers were globally resolved to 9.3 Å and 9.6 Å, respectively (Fig. 2A, 2B, Fig. S3, Table S2), with local resolution ranging from 8 Å – 16 Å (Fig. S3). At this resolution, Arf1 and Nef could be unambiguously fit to the density. Two Arf1 and one Nef molecules per AP-1 complex constitute the asymmetric unit of the lattice (Fig. 2C). The stoichiometry and interfaces between AP-1, Arf1 and Nef within the asymmetric unit match those observed in the asymmetric unit of the soluble (e.g. non-membrane associated) closed trimeric AP-1 complex consisting of the AP-1^core^, N-terminally truncated Arf1 and a genetic fusion of BST2^(2-20)^ and NL4-3 Nef (PDB: 6cm9) (Morris et al., 2018). Indeed, the asymmetric unit of the closed trimer docks into the lattice density with no need for conformational adjustments, and only slight shift in the Nef core. The major interactions between the β1-bound Arf1 (hereafter, Arf1^β1^) γ-bound Arf1 (hereafter, Arf1^γ^) with the GTP-dependent switch regions of the Arf1 molecules is identical to the seen in the BST2^(2-20)^-bound closed AP-1 trimer (Morris et al., 2018), and therefore not presented in detail here. The density of the lattice is fully accounted for by AP-1^core^, Arf1^myr^ and Nef^myr^.

**Figure 2.**
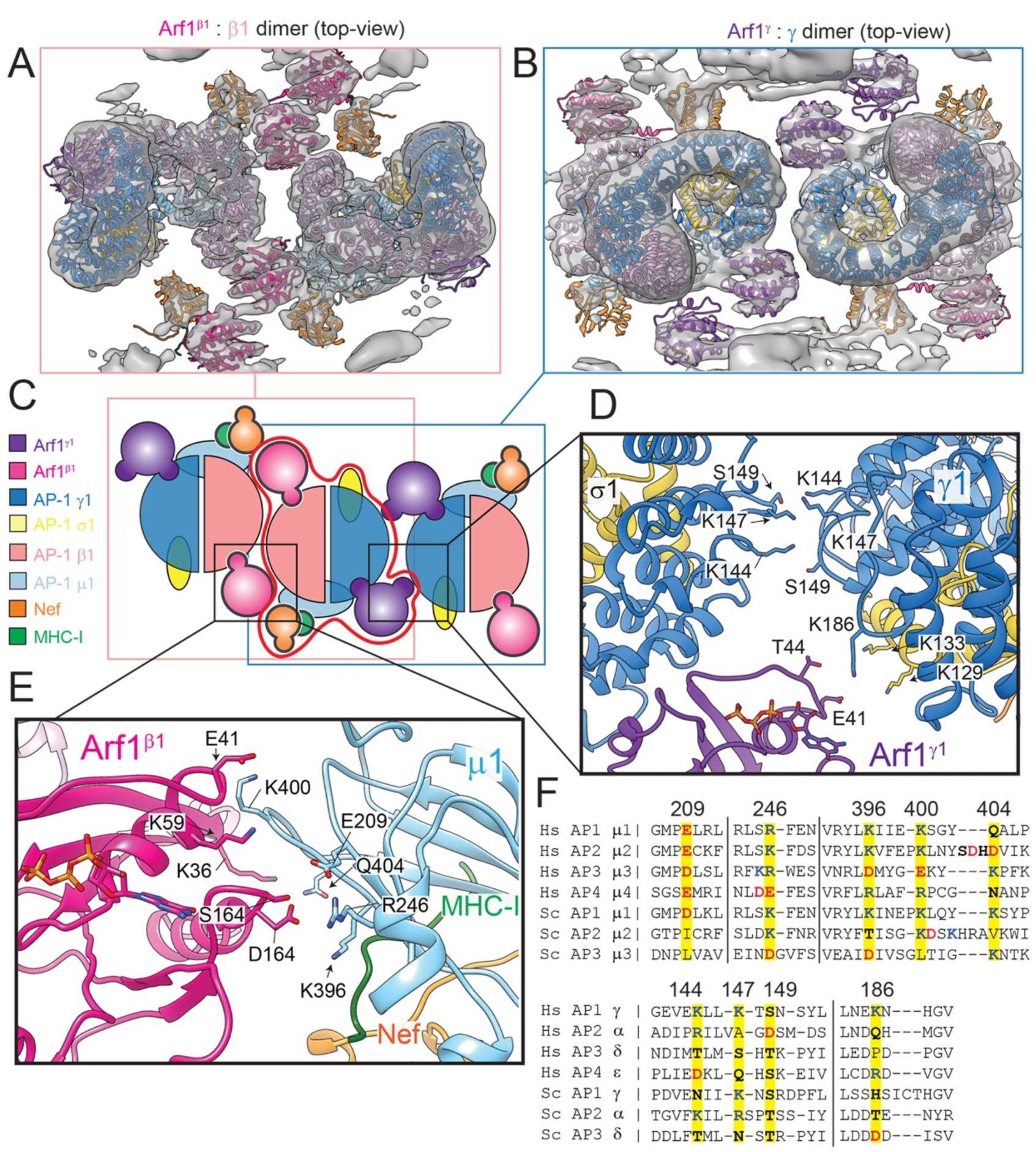
Arf1 mediates AP-1 dimerization. (A) Sub-nanometer resolution map and associated atomic model fit of the Arf1^β1^ : β1 juxtaposed AP-1 dimer as viewed looking onto the tube surface. Atomic models are colored as denoted in (C). (B) Sub-nanometer resolution map and associated atomic model fit of the Arf1^γ^ : γ juxtaposed AP-1 dimer as viewed looking onto the tube surface. Atomic models are colored as denoted in (C). (C) Cartoon representation of the asymmetric unit (AP-1_1_ : Arf1_2_ : Nef_1_; red outline) flanked on either side by asymmetric units creating Arf1^γ^ : γ and Arf1^β1^ : β1 dimeric connections. (D) Zoom of the interface mediating Arf1^γ^ : γ and γ : γ dimeric interactions. (E) Zoom of the interface mediating Arf1^β1^ : β1 dimeric interactions. (F) Sequence alignment of μ and γ (or equivalent) domains from human and yeast APs. Yellow highlighted residues are labeled according to their position in human AP-1 and correlate with residues shown in panels (D) and (E).

Placement of the asymmetric unit (AP-1_1_:Arf1_2_:Nef_1_) within each dimer revealed that the AP-1 protomers do not contact one another directly. Rather, each dimeric interface is mediated by AP-1:Arf1 and Arf1:Arf1 contacts. These Arf1 dimer contacts have not been reported previously. At 9 Å resolution, side-chain densities are not visible, however, side-chain locations can be inferred from the previously determined high-resolution structures of all of the components. Arf1^β1^ localizes charged and polar residues (K36, E41, K59, S162 and D164) near complementary residues in the μ1 domain of an adjacent asymmetric unit (R246, K396, K400, Q404) (Fig. 2E). Arf1^γ^ similarly localizes E41 and T44 with γ K186 and σ1 K129/K133 (Fig. 2D). Arf1 switch 1 and 2 regions consist of residues 45-54 and 70-80 respectively. Thus, some of these contacts are at the periphery of the GTP-dependent conformational switch regions, but none are actually within the switch regions. The biochemical properties of residues at each interface, particularly the Arf1-μ1 interface, are conserved between AP-1 and its human and yeast AP homologs (Fig. 2F). This suggests that the other Arf1-dependent AP complexes could use similar contacts to form Arf1-mediated higher order assemblies. Consistent with this, an Arf1 contact with the CTD of the μ4 subunit of AP-4 was inferred from a yeast two-hybrid binding screen (Boehm et al., 2001). The only direct AP-1:AP-1 interaction found within the lattice is created by γ residues K144, K147 and S149 (Fig. 2E). AP-2, -3 and -4 also share polar and charged residues at these positions (Fig. 2F), again suggesting other APs could make similar contacts.

Next, we analyzed interactions that stabilize axial propagation of AP-1 protomers along the tube axis. The two respective axial interfaces bridging β1:β1 and γ:γ dimers were symmetry expanded and the STA maps globally refined to 9.3 Å and 9.4 Å, respectively (Fig. 3A, B, Fig. S3, Table S2). Each interface is primarily composed of direct Arf1:Arf1 interactions and both interfaces utilize a core set of residues conserved amongst human Arf1 homologs and yeast Arf1 (Fig. 3E), namely E115, R117, R149 and H150 (Fig. 3C, D). Arf1^γ^ also makes polar contacts between its H146 and μ1 E284 of the adjacent AP-1 protomer Fig. 3C.

**Figure 3.**
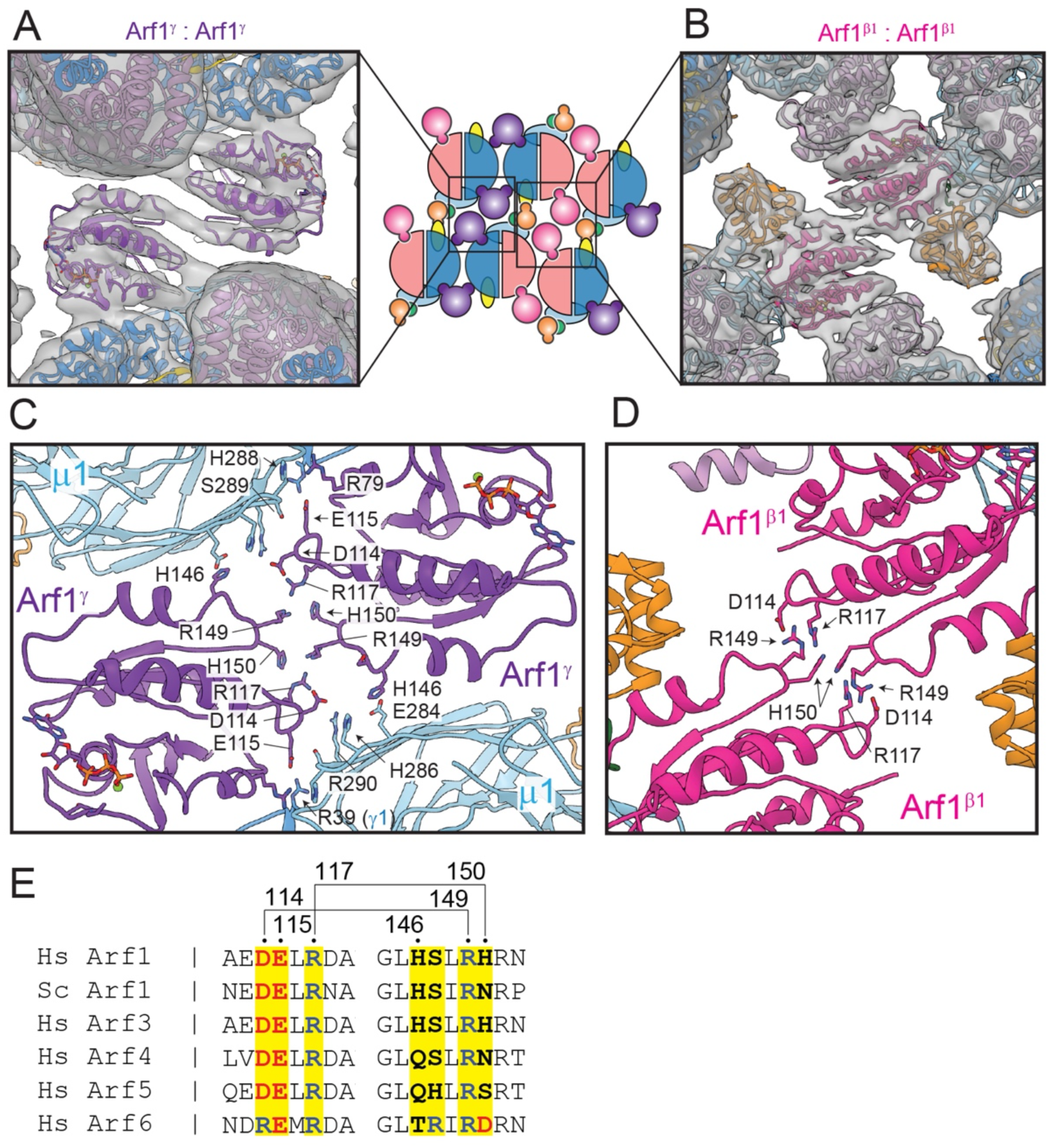
Arf1:Arf1 homodimerization in the narrow tube lattice. (A) Sub-nanometer resolution map and associated atomic model fit of the Arf1^γ^ : Arf1^γ^ homodimeric interface as viewed looking onto the tube surface. Atomic models are colored as denoted in Fig. 2. Cartoon representation of the AP-1 lattice as viewed looking onto the tube surface is shown for reference. (B) Sub-nanometer resolution map and associated atomic model fit of the Arf1^β1^ : Arf1^β1^ homodimeric interface as viewed looking onto the tube surface. Atomic models are colored as denoted in Fig. 2. Cartoon representation of the AP-1 lattice as viewed looking onto the tube surface is shown for reference.(C) Zoom of the interface mediating Arf1^γ^ : Arf1^γ^ homodimerization. (D) Zoom of the interface mediating Arf1^β1^ : Arf1^β1^ homodimerization. (E) Sequence alignment of Arf1 (human and yeast) and human Arf1 homologs. Yellow highlighted residues are labeled according to their position in human Arf1 and correlate with residues shown in panels (C) and (D). Residues linked by solid lines are proposed to interact directly.

### Arf1, Nef and AP-1 contributions to membrane binding

Each protomer is arranged on the membrane surface such that the N-termini of Arf1 and Nef are proximal to and oriented towards the membrane (Fig. 4A, B). Density for the N-terminal residues of Arf1 (2-17) is resolved well enough to model an amphipathic helix and unstructured chain leading to the Arf1 globular core (Fig. 4C, D). The modeled amphipathic helix is oriented along the long axis of the membrane tube consistent with the map density (Fig. 4C-E), although the relative orientation is more ambiguous for Arf1^β1^ (Fig. 4D,E). The corresponding density for the N-terminus of Nef is not resolved in any of the sub-nanometer resolution structures, including those with no symmetry imposed during subtomogram alignment, suggesting the position and/or orientation of its N-terminus is less constrained relative to the N-terminus of Arf1.

**Figure 4.**
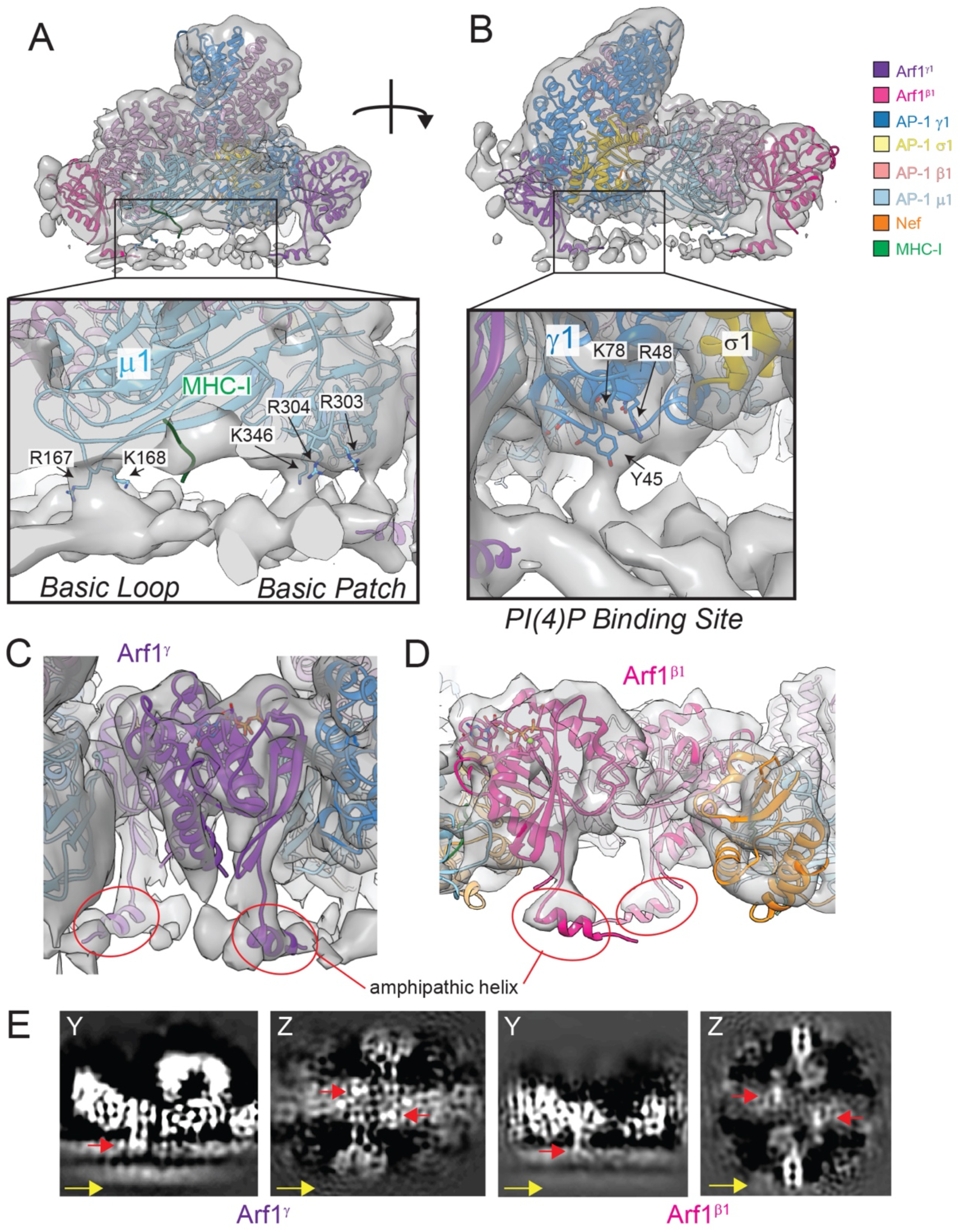
Lattice contacts with the membrane. (A) Sub-nanometer resolution map and associated atomic model fit of the narrow tube lattice asymmetric unit (AP-1_1_ : Arf1_2_ : Nef_1_) as viewed from the side. Atomic models are colored as denoted in the legend shown in panel (B). The inset shows a zoom of the AP-1 : membrane interface specifically highlighting μ1 : membrane contacts. Nef is omitted for visual clarity. (B) 180° rotated view the map and model shown in (A). The inset shows a zoom of the AP- 1 : membrane interface specifically highlighting γ : membrane contacts. (C) Atomic model fit to the density associated with Arf1^γ^ as viewed from the side. Density for the near-complete N-terminus (2-17) is present, and a speculative model for the amphipathic helix is shown and highlighted (red circles). (D) Atomic model fit to the density associated with Arf1^β1^ as viewed from the side. As in (C), density for the near-complete N-terminus (2-17) is present, and a speculative model for the amphipathic helix is shown and highlighted (red circles). (E) Y and Z slices through the STA structure(s) at regions of Arf1 : membrane contacts. Slices emphasize the density associated with the Arf1 N-terminus. The Y slice shows Arf1 density from the side, while the Z slice shows Arf1 density in-plane with the membrane. Red arrows denote density assigned to the putative Arf1 amphipathic helix. The yellow arrow denotes the long axis of the tube.

The AP-1 core is oriented such that μ1 and γ are parallel with the membrane and make primarily three points of contact. Residues R167 and K168 within the basic loop of μ1 and K346 within the basic patch of μ1 appear to make direct contact with the membrane, presumably with anionic lipid headgroups. R303 and R304 in the basic patch are also closely localized to the membrane surface (Fig. 4A). The third point of contact is observed in the γ N-terminus (Fig. 4B). Residues predicted to interact with phosphoinositides, e.g. PI(4)P, including Y45, K78 and R48 are localized to membrane density near the putative PIP headgroup binding site, suggesting AP-1 γ is engaged with PI(4)P (Heldwein et al., 2004).

### Nef and MHC-I engagement with the tubular coat

Rigid body docking of the AP-1:Arf1_2_:BST2-Nef structure (PDB: 6CM9) confirms Nef is localized immediately adjacent to the AP-1 μ1 tyrosine motif binding site within the AP- 1:Arf1 tubular lattice (Fig. 2A, 3B, 5C). The crystalized Nef, μ1 and MHC-I structure (PDB: 4EN2, (Jia et al., 2012)) can be docked into the STA essentially without modification (Fig. 5C, 5D). The Nef 72-PxxPxxP-78 motif is positioned to complement the MHC-I^ctyo^ tail at the μ1 cargo binding site and the Nef N-terminal helix (9-24) is packed against the Nef core (Fig. 5D). Residues upstream of the Nef N-terminal helix are proximal to the membrane and likely directly interact with the membrane in a manner consistent with NMR structures of the N-terminal membrane anchor domain of Nef (Grzesiek et al., 1996). As seen in the Nef:MHC-I:μ1 crystal structure (PDB: 4EN2, (Jia et al., 2012)) and AP-1:Arf1:BST2-Nef complex (Morris et al., 2018), the acidic patch of Nef (62-EEEE-65) is only partially resolved in the STA density, and direct interaction can only be inferred between μ1 K302 and Nef E65. The entire loop spanning from the end of the N-terminal helix to the acidic patch (E24-E64) is unresolved (Fig. 5D). Overall, our observations in the context of the lipid bilayer are consistent with the direct interactions between Nef, μ1 and the cytosolic tails of MHC-I and tetherin seen previously in the absence of membrane (Fig. 5C, 5D) (Jia et al., 2012; Morris et al., 2018).

**Figure 5.**
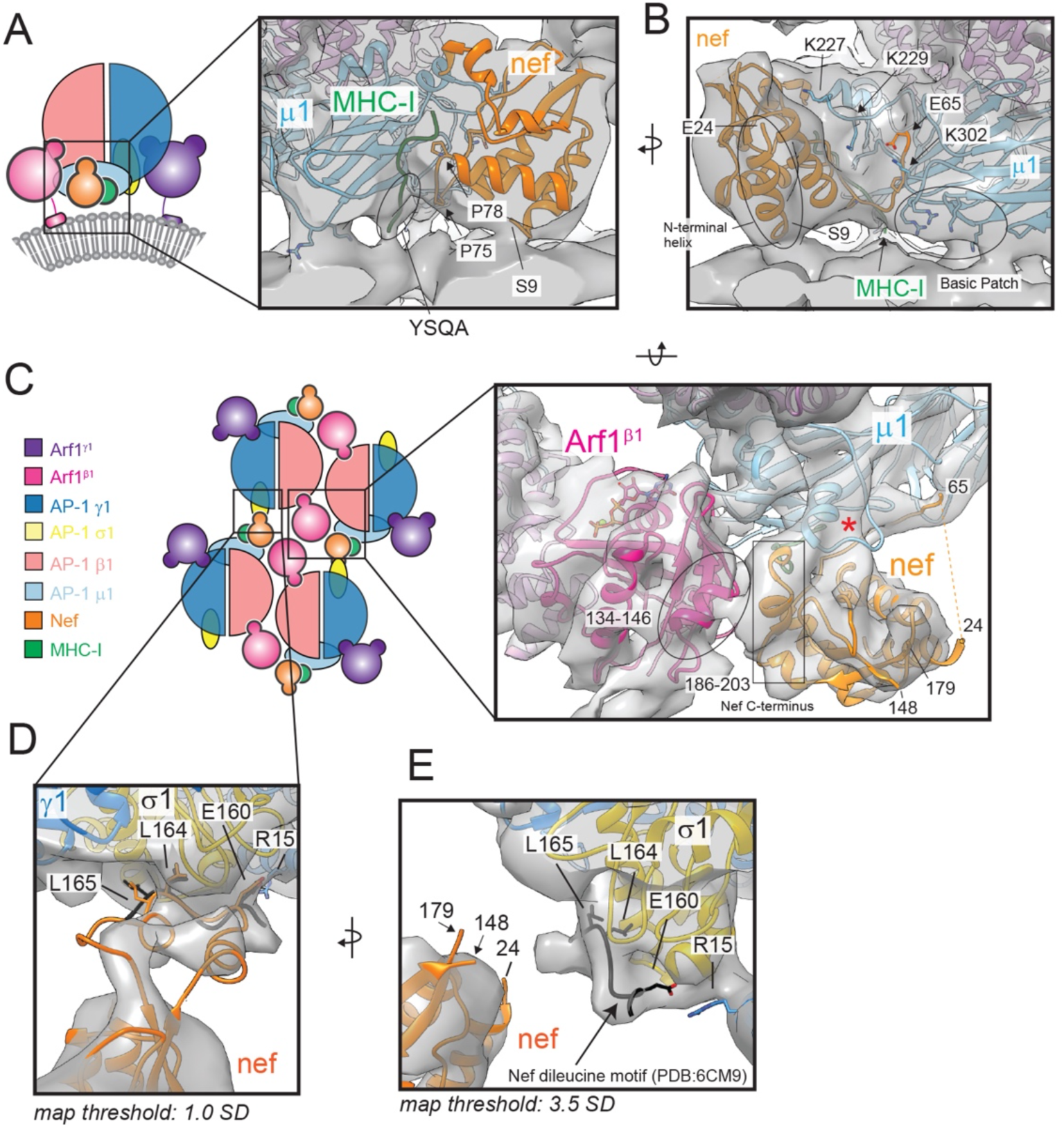
Nef interactions within the AP-1:Ar1 lattice. (A) Side-view cartoon depiction of the AP-1:Arf1_2_:Nef asymmetric unit from the narrow tube lattice with inset showing the Nef : MHC-I : μ1 interfaces. (B) 180° rotated view of the inset shown in (A). (C) Top- view cartoon depiction of the AP-1:Arf1:Nef lattice with inset showing the Nef : μ1 : MHC-I : Arf1^β1^ quaternary complex (map threshold: 3.5 SD). The red asterisk denotes the μ1 internal loop stabilized by Nef. (D) Inset showing the Nef ExxxLL : σ1 interface. The map density is fit with a hypothetical structural model based on predicted and solved structures. The ExxxLL motif of Nef observed in the AP-1:Arf1:BST2-Nef closed trimer (PDB:6CM9) is shown in black for comparison. (E) 90° rotated view of (D). The map threshold density was set to 3.5 SD from the mean. The ExxxLL motif of Nef from PDB: 6cm9 (black) is modeled as in (D). The Y slice shows Arf1 density from the side, while the Z slice shows Arf1 density in-plane with the membrane. Red arrows denote density assigned to the putative Arf1 amphipathic helix. The yellow arrow denotes the long axis of the tube.

Nef makes two additional interactions in the lattice context that have not been previously reported. Directly adjacent to the tyrosine cargo binding site, the C-terminal residues 186-203 pack against a helical region of Arf1^β1^ consisting of residues 134-146 (Fig. 5C). The same C-terminal region of Nef stabilizes an internal loop on μ1 and facilitates ternary contacts between MHC-I, Nef and μ1 (Fig. 5C) as previously observed (Jia et al., 2012)). The proximity of Arf1^β1^ to the cargo binding groove, at a region distinct from the YSQA binding region, hints at a potential role for Arf1 in cargo binding and/or cargo specificity, especially for cargoes with longer cytosolic tails. The other noteworthy interaction between Nef and AP-1 is mediated by the Nef dileucine motif (160-ExxxLL- 165) (Fig. 5A, 5B). Nef is positioned within the lattice such that its dileucine motif reaches across to an adjacent AP-1 protomer (Fig. 5A, 5B). The dileucine site of σ1 is clearly occupied (Fig. 5B) consistent with the σ1:160-ExxxLL-165 interaction observed in the AP- 1:Arf1:BST-2 closed trimer (Morris et al., 2018) and its known interaction with AP-1 (Bresnahan et al., 1998; Janvier et al., 2003a; Janvier et al., 2003b). Increasing the STA map threshold revealed density directly linking the σ1 site to Nef’s core (Fig. 5A). Docking of predicted and previously solved structures of Nef’s dileucine loop into the map correlate well with the density and meet the spatial requirements for a direct interaction (Fig. 5A), thus supporting Nef interactions with two distinct AP-1 molecules in the lattice.

### Rearrangements stabilize the lattice on wide tubes

Ordered AP-1 coats were observed on the majority of membrane tubes, including those with much wider diameters (e.g. >60nm, Fig. S4A). To determine how the lattice adjusts to fit flatter membrane surfaces, subtomograms from wider tubes (60-70nm diameter) were aligned and averaged. The resulting maps were globally resolved to 20 Å (Fig. S4B, Table S3). The helical pitch on wide tubes is 70 nm (Table S4) and AP-1 dimers are stacked almost coincident with the long axis of the membrane tube (Fig. 1C, 1D vs Fig. S4A, S4F). As on narrow tubes, the lattice is stabilized by AP-1:Arf1 and Arf1:Arf1 dimeric interactions, and asymmetric units within the lattice can be fit with the same AP- 1:Arf1_2_:Nef promoter derived from the closed, trimeric AP-1 assembly (Fig. 6A), with notable exceptions, discussed below. Interactions with the membrane are conserved between the narrow and wide tube lattices with clear and obvious density for the N-termini of Arf1 extending downward from the protein coat to the outer leaflet of the bilayer (Fig. 6B, S4E), and contacts between the membrane and AP-1 subunits (Fig. S4C, S4E).

**Figure 6.**
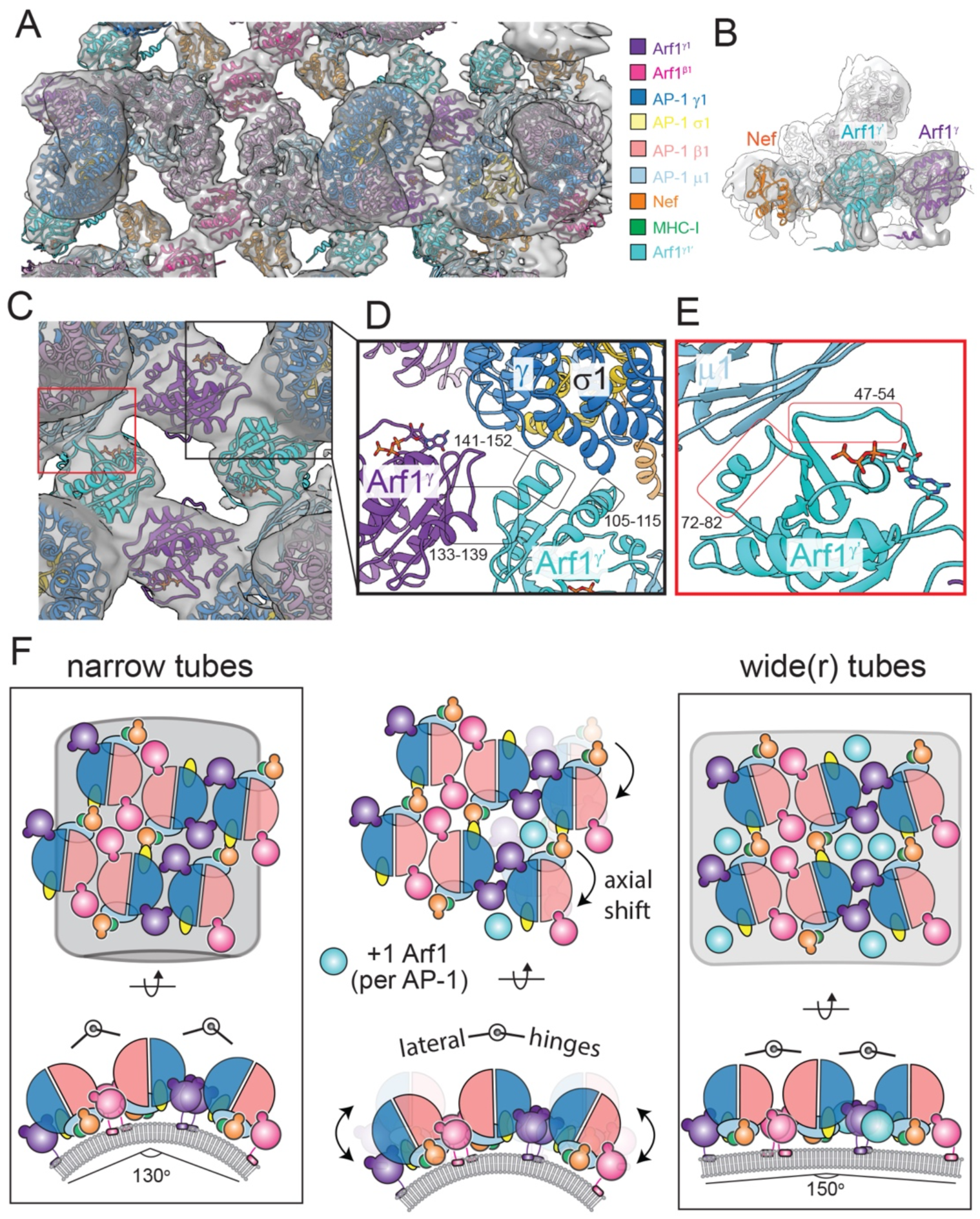
Rearrangements of the lattice on wide tubes. (A) Composite map and atomic model of the Arf1^β1^-linked AP-1 dimer and Arf1^γ^-linked dimer on wide tubes, shown from the top-view. Arf1^γ’^, the third Arf1 molecule within an asymmetric unit on wide tubes, is shown in cyan. (B) Side view of the STA map with atomic model fit of Arf1^γ’^. (C) Composite map and atomic model of the Arf1^γ^ axial interface, shown from the top-view. (D) Zoom of the Arf1^γ^ - Arf1^γ’^ homodimeric interface, Arf1^γ^ : γ interface and Arf1^γ’^ : γ/σ1 interface. (E) Zoom of the Arf1^γ’^ : μ1 interface. (F) Cartoon summary of the AP- 1:Arf1:Nef lattice interactions on narrow and wide tubes. The left and right cartoons depict the arrangements structurally observed on tubes. The middle cartoon illustrates the differences between the two arrangements.

Symmetry expansion and focused alignment and averaging of the asymmetric unit and individual dimeric interfaces resulted in EM density maps globally resolved to 9.2 Å for the asymmetric unit, 9.8 Å for the β1:β1-juxtaposed dimer and 11.6 Å for the γ:γ- juxtaposed AP-1 dimer (Fig. S4C, Table S2). At the γ:Arf1 dimeric interface, the γ subunits are no longer close enough to directly interact, due to a ∼3 nm displacement of one AP- 1 protomer along the tube axis (Fig. S4E). Interactions between Arf1^γ^ and the juxtaposed γ subunit stabilize the new dimeric interactions (Fig. 6D). The relative angle between asymmetric units at the γ:Arf1 dimeric interface increases from 130° on narrow tubes to 150° on wide tubes, but without involving significant changes to the asymmetric unit itself. The β1:Arf1 dimeric interface is similar to the interface on narrow tubes, with the exception that the interface is straighter by 20° (Fig. 6F, S4B). The asymmetric units appear to hinge at the Arf1^β1^:β1 interface. With the new arrangement, new polar and electrostatic interactions are mediated between Arf1^β1^ D164/Y167 and μ1 R246/K400 (Fig. 6D, S4C). At the homodimeric Arf1^β1^ interface, the contacting residues appear to be the same as those in the narrow lattice indicating the interactions are conserved between narrow and wide tube lattices. Contacts between Nef, cargo, AP-1 and the membrane are also unchanged.

The most prominent deviation between wide and narrow lattices is the presence of additional protein density at the axial interface between Arf1^γ^:γ dimer stacks (Fig. 6C vs Fig. 3B) belonging to an additional Arf1 molecule. Due to the local resolution of the maps (10-12Å; Fig. S4D), rigid-body modeling of Arf1 resulted in multiple Arf1 arrangements that could satisfactorily fit the density. One solution is shown for clarity (Fig. 6A-E). The result reveals that 3 Arf1 and 1 Nef per AP-1 constitutes the asymmetric unit within the lattice on wide tubes. The auxiliary Arf1, termed Arf1^γ’^, is positioned to simultaneously contact μ1, γ and Arf1^γ^ (Fig. 6A-C, 6F). The interfacial contacts map to the same general locations as those seen in the narrow tube lattice (Fig. 3B, 6F). The one exception is that Arf1^γ’^ creates a novel homotypic interface with Arf1^γ^ using residues 133- 139 which are distinct from Arf1:Arf1 dimer contacts in the narrow lattice. On the basis of the novel Arf1 dimers observed on membranes in this study, we revisited our previously predicted model for a hexagonal AP-1:Arf1 coat compatible with a hexagonal clathrin lattice (Shen et al., 2015). In the new model (Fig. S5), the flexibility of the Arf1 contacts observed here gives rise to a mixture of pentagons and hexagons, instead of hexagons only, and is thus fully compatible with formation of a dodecahedral clathrin cage. This mixture of pentagons and hexagons results in a model for a lower density spherical coat forming the inner layer of AP-1 clathrin vesicles, whose character thus differs dramatically from the clathrin-independent tubular coat structure reported here.

### AP-1 tubes are stabilized by clathrin knockdown and colocalize with Nef in cells

In cells, AP-1 and Arf1 localize to tubular and vesicular membrane compartments, such as the TGN, endosomes, and clathrin coated vesicles, during normal cargo trafficking (Futter et al., 1998; Huang et al., 2001; Robinson, 1990; Stamnes and Rothman, 1993). Nef expression promotes AP-1 localization to tubulated membranes (Janvier et al., 2003a; Lubben et al., 2007). As detailed above, the AP-1 lattice is stabilized by multivalent contacts. Arf1:AP-1 interactions mediate both axial and lateral contacts within the lattice, suggesting Nef augments but is not essential for lattice formation. Thus, we hypothesized that AP-1 and Arf1 could self-assemble into tubular coats in the absence of Nef, and that Nef would localize to these structures in cells. Unlike AP-3 and -4, nearly all reported AP- 1-dependent processes are clathrin-dependent. We thus further reasoned that the tubular clathrin-independent AP-1 coat would represent a clathrin-independent intermediate preceding the formation of CCVs, not a fully clathrin-independent pathway. This hypothesis predicts that clathrin knockdown should stabilize such an intermediate, enriching the cell with AP-1 positive tubules.

AP-1 and Nef localization was monitored in HeLa cells by Airyscan confocal microscopy. AP-1 complexes were visualized by expression a μ1A construct C-terminally tagged with GFP (Guo et al., 2013) and Nef was C-terminally HALO-tagged. In the absence of Nef, AP-1 localized to spherical densities throughout the cytosol and at the perinuclear region (Fig. 7A). In the presence of Nef, the number and morphology of AP- 1 densities remained virtually unchanged (Fig. 7B). Nef signals partially overlapped with AP-1 density under these conditions (Fig 7D). Notably, the AP-1 signal more strongly overlapped with the perinuclear region in Nef expressing cells (compare Fig. 7A vs 7B).

**Figure 7.**
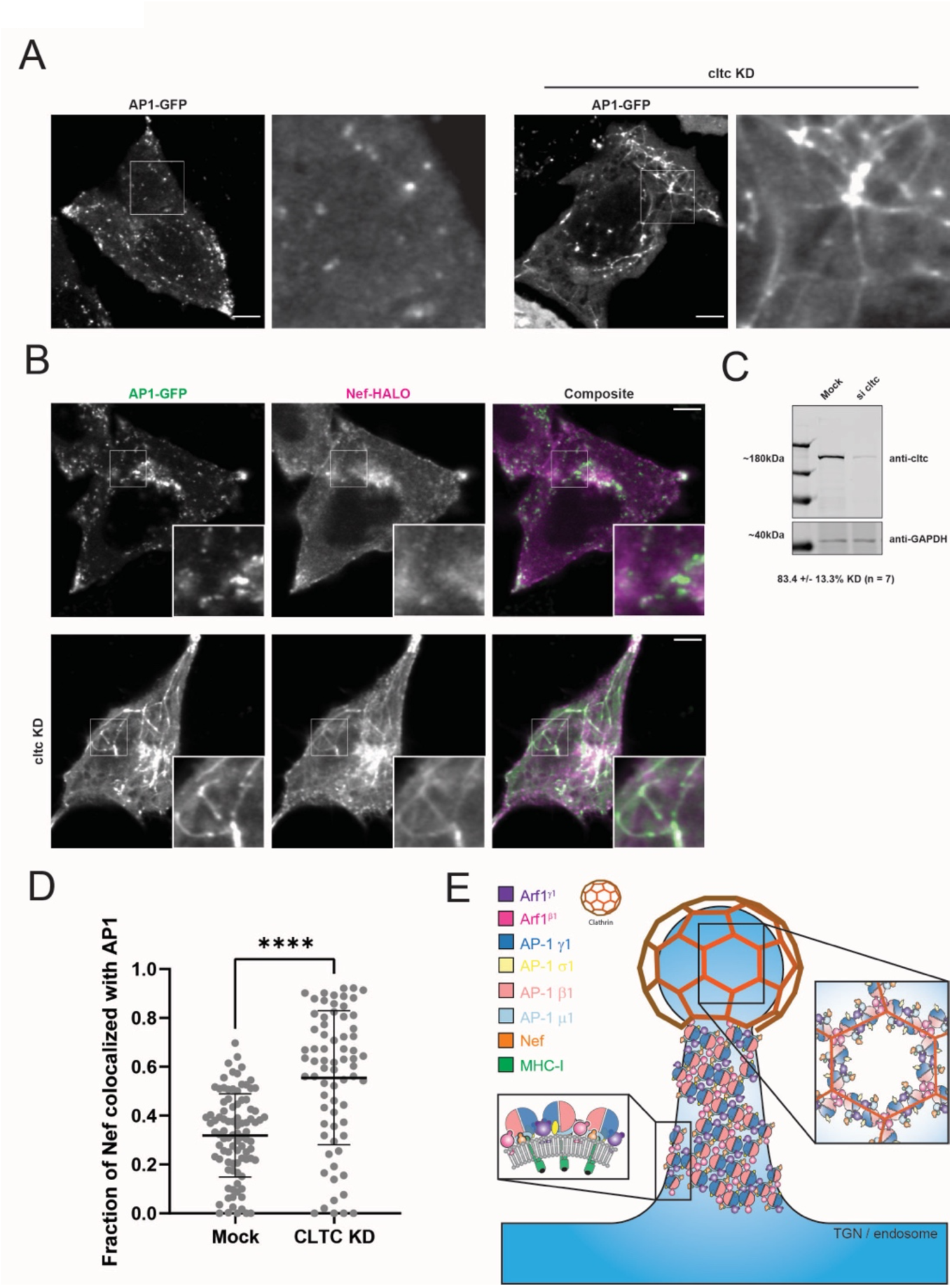
Clathrin knockdown induces formation of AP-1 tubules in cellulo.(A) Single Z-slice images of HeLa cells transfected with AP-1-GFP construct under mock or clathrin knockdown conditions, acquired using live cell confocal microscopy. Magnified insets of the indicated area are shown on the right of each image. Scale bar, 5µm. (B) Single Z-slice images of HeLa cells cotransfected with AP-1 GFP and NA7-Nef-HALO construct under mock or clathrin knockdown conditions, acquired using live cell confocal microscopy. Magnified insets of the indicated area are shown on the lower right corner of image. White arrows point to spheroid structures where AP1 and Nef colocalize.Scale bar, 5µm. (C) Western blot of mock or clathrin knockdown (si cltc) HeLa cell lysate 96 hours after siRNA treatment, probed with anti-cltc and anti-GAPDH antibodies. Average % of cltc knockdown +/- SD shown at bottom. (D) Quantification of AP-1 and Nef colocalization in the presence and absence of clathrin. Line indicates mean. (E) Model. Multimodal interactions account for versatile arrangements of AP-1 and Arf1 on membranes. Lattice like AP-1:Arf1 assemblies on membrane tubes accumulate cargo in trafficking compartments. Hexagonal and pentagonal AP-1:Arf1 assemblies interface with clathrin to mobilize cargo into vesicles. Nef hijacks the AP-1:Arf1 tubular coat to sequester cargo by promoting lattice arrangements and co-opts the clathrin-like geometries to promote trafficking of cargoes to the lysosome.

We then replicated the observation that when clathrin is depleted in cells, AP-1 is enriched on tubular endosomes (Janvier and Bonifacino, 2005). Clathrin was depleted from cells by siRNA and the localization of AP-1 and Nef was monitored. Under knockdown conditions, clathrin heavy chain levels were reduced on average by >80% as determined by western blot (Fig. 7C). AP-1 robustly localized to tubulated membranes. Clathrin knockdown increased Nef association with both punctate and tubular AP-1 structures in a statistically significant manner (Fig. 7D). This suggests that Nef preferentially associated with the tubular form of AP-1 coat that predominates in clathrin knockdown. These observations confirm that membrane tubulation is an inherent property of AP-1 that is enhanced by the removal of clathrin. These AP-1 tubules strongly co- localize with Nef when it is expressed, yet their formation does not require Nef.

## Discussion

The Arf-dependent APs have long been observed on tubular structures in cells and possess clathrin-independent functions, yet the mechanism of their assembly on tubulovesicular membranes has been unknown. It was not even clear if tubular coat was an inherent property of AP complexes or required additional factors. Here, we took advantage of the ability of the HIV-1 Nef protein to overdrive AP-1 tubular vesicle formation in MHC-I downregulation. We confirmed that tubular membrane coat formation is an inherent property of AP-1, requiring only Arf1 as a cofactor. We went on the determine the structure of the coat. The coat is densely packed on membranes, similar in that respect to retromer (Kovtun et al., 2018), leaving few gaps on the membrane surface.

AP-1:Arf strongly promotes formation of high curvature tubules (Fig. S1E), although it can assemble on flat membranes as well. There is an extensive literature showing that Arf1 associates with tubulated membranes and can, at sufficiently high concentrations, stabilize curvature and induce tubulation by itself (Beck et al., 2011; Beck et al., 2008; Bottanelli et al., 2017; Diestelkoetter-Bachert et al., 2020; Krauss et al., 2008). Isolated Arf1 has been previously shown to dimerize (Diestelkoetter-Bachert et al., 2020), but not to former higher assemblies on its own. Curvature induction by Arf1 *in vivo* therefore requires a mechanism to cluster Arf1 at the needed high local concentration. AP-1 recruitment is intimately linked to Arf1 (Stamnes and Rothman, 1993; Traub et al., 1993). AP-1 has multiple Arf1 binding sites (Morris et al., 2018; Ren et al., 2013), which provides a means to cross-link Arf1 into larger assemblies. On the basis of their known Arf1 dependence and structural conservation, the same can be said for AP-3 (Ooi et al., 1998) and AP-4 (Boehm et al., 2001). Thus, cellular Arf1-localized tubules (Bottanelli et al., 2017) most likely consist of Arf1 dimers cross-linked by AP complexes.

The structural model of the AP-1 membrane coat was built on a body of prior crystallographic and cryo-EM analysis in the absence of membranes. Some features are as anticipated from past work, while others are not. AP-1 protomer docking on membranes is essentially as predicted (Jia et al., 2014; Ren et al., 2013; Shen et al., 2015). The protomer docking mode is also consistent with the docking seen in reconstructions of the AP-2 complex within clathrin-coated vesicles (Kovtun et al., 2020; Paraan et al., 2020). The lattice observed here is more densely packed than that of the open hexagonal AP-1:Arf1 lattice proposed to recruit clathrin (Shen et al., 2015) (Fig. 7G, S5), and is incompatible with clathrin binding. Previously, the structure of a membrane-free assembly of a closed AP-1:Arf1:Nef:BST2 trimer was determined (Morris et al., 2018), in which Arf1^β^, but not Arf1γ, were disposed on the membrane-binding face. In the presence of membranes and the MHC-I tail, we find here that both Arf1^β^ and Arf1^γ^ contact membranes. The closed AP-1:Arf1:Nef:BST2 trimer (Morris et al., 2018) was also incompatible with propagation into a higher order lattice. The fundamental biological insight here is that AP-1:Arf1 can form a dense coat and thereby tubulate membranes on its own, in the absence of clathrin. The tubular structure is denser than the clathrin-associated version of the AP-1:Arf1 assembly. The high density is made possible by novel Arf1 dimer-mediated bridging, which allows for tighter AP-1 packing than is possible with the previously known Arf1 trimers.

These new data and models put the role of Nef in MHC-I sequestration and degradation into fresh perspective. The structure shows that Arf1 cross-links AP-1 laterally and axially and is sufficient on its own to bridge the tubular coat. In cells, we observe that clathrin knockdown is sufficient to induce extensive AP-1 tubular structures, whether or not Nef is present. Once present, Nef strongly colocalizes with the tubes. Nef integration into the tubules provides a mechanism for sequestration of MHC-I (Blagoveshchenskaya et al., 2002; Dirk et al., 2016; Janvier et al., 2003a; Le Gall et al., 1998; Tavares et al., 2020). We observed that the Nef dileucine motif is positioned to cross-link adjacent AP-1 protomers in the tubular lattice. Nef does not require its dileucine motif to downregulate MHC-I (Greenberg et al., 1998; Mangasarian et al., 1999; Roeth et al., 2004). The interactions between μ1, the Nef core, and the YQSA motif of MHC-I are sufficient for downregulation from the plasma membrane without needing the dileucine motif. However, the Nef dileucine motif does increase the time that AP-1 (and AP-3) complexes reside on endosomal membranes (Coleman et al., 2006; Janvier et al., 2003a). This effect is consistent with the observation that Nef colocalization with AP-1 increases upon clathrin knockdown, which favors the tubular AP-1 lattice over the open clathrin-associated lattice.

Arf1 trimers were previously observed in the context of AP-1 in the absence of membranes (Shen et al., 2015; Morris et al., 2018) and shown to form a hexagonal lattice on membranes (Shen et al., 2015). Arf1 trimer-linked COPI coats have been directly visualized on spherical membrane vesicles (Dodonova et al., 2017). The different geometries and interfaces formed by Arf1 in different settings suggests the energy barriers between the different arrangements must be small. The interfacial free energy of association of each of the molecular contacts is also likely to be small, given that they are dominated by polar residues and bury modest amounts of solvent accessible surface area. We generated a model for the AP-1:Arf1 lattice rearrangements that would match the symmetry of a clathrin coat (Fig. 7G, S5). Given clathrin depletion stabilizes AP-1 tubule formation, we infer that the dense tubular coat form of AP-1 exists normally in the presence of clathrin and absence of Nef, but as a transient intermediate in CCV formation. The rearrangement to a clathrin-like geometry entails breaking one set of dimeric contacts and replacing them with trimeric contacts. The polar character of the dimer contacts helps ensure these contacts are weak and can break readily when a rearrangement is needed. The idea that a dense cylindrical AP-1:Arf1 inner coat could exist as a normal intermediate in CCV formation is consistent with new thinking about the role of the COPII inner coat at ER exit sites (Melero et al., 2022). In this model, the inner coat sets the radius of membrane curvature, and initiates the subsequent recruitment of clathrin. Once clathrin begins to arrive, AP-1:Arf1 rearranges into hexagons and pentagons (Fig. 7G, S5), leading to CCV formation.

Our finding that the Arf1 interaction sites are largely conserved across species and between AP-1, -3, and -4 suggest our observations are general across biology and span disparate pathways mediated by the Arf1-dependent AP complexes. We proposed above that the AP-1:Arf1 tubular coat ordinary functions as an intermediate on the pathway to CCV assembly. This idea is supported by the increased AP-1 tubule formation upon clathrin knockdown. We proposed that HIV-1 Nef hijacks this tubular intermediate and causes it to persist for an abnormally time to sequester MHC-I, prior to its eventual packaging in CCVs and sorting to the lysosome for degradation. Since AP-3 and AP-4 activities are largely (Peden et al., 2004; Vowels and Payne, 1998) or completely (Dell’Angelica et al., 1999; Hirst et al., 1999) independent of clathrin, we predict that these two complexes self-assemble with Arf1 into stable tubular coats. For AP-3 and -4, these tubular entities would presumably be the cargo carriers themselves, not merely intermediates in spherical vesicle formation. While the AP-1 tubule would be terminated by the recruitment of clathrin, the AP-3 and AP-4 tubules should persist until GTP hydrolysis, catalyzed by Arf GAPs, promotes dissociation of Arf1 from the membrane.

In summary, we found that AP-1:Arf1 can form a contiguous tubular membrane coat cross-linked by Arf1 dimers, even in the absence of clathrin, and we determined the structure of the coat. This has allowed us to structurally rationalize a large body of unexplained findings on the clathrin-independent roles of AP complexes, their residence on tubular structures, and the ability of AP-1 to sequester MHC-I during Nef-dependent downregulation.

## Acknowledgments

We thank J. Hutchings and G. Zanetti for providing a data set for workflow validation, D. Toso and J. Remis for cryo-EM operational support, and D. Drubin for use of the Zeiss LSM900 microscope. This research was supported by NIH grants F32 AI152971 (R. H.), R01 AI120691 (J.H.H.), and P50 AI150476 (J.H.H.).

## Methods and Materials

### DNA constructs for protein expression

The His6- and GST-tagged AP-1 constructs were previously described (Shen et al., 2015). For the full-length beta 1 AP-1 (AP-1^βFL^) complex, β1 (1-584) was replaced with full-length β1 DNA. MHC-I (338-365) peptide was expressed as a TEV-cleavable C-terminal fusion to GST and included a C-terminal His6 tag and N-terminal cysteine for lipopeptide coupling. NL4-3 Nef was expressed with a TEV-cleavable C-terminal His6 tag from the pST39 expression vector. Yeast NMT or Human NMT1 (81-496) was subcloned into the pRSFDuet-1 vector. Bovine Arf1 (no tag) was subcloned into the pHis-parallel2 vector using NdeI/XhoI sites. The bovine Arf1 protein sequence is the same as human.

### Protein purification

The AP-1 complexes were expressed in BL21 (DE3) pLysS (Promega, Madison, WI) strains and induced with 0.3 mM isopropyl-β-D-thiogalactopyranoside (IPTG) at 20°C overnight. The cells were lysed by sonication in 50 mM Tris at pH 8.0, 300 mM NaCl, 10% glycerol, 3 mM β-mercaptoethanol (β-ME), and 0.5 mM phenylmethanesulfonyl fluoride (PMSF). The clarified lysate was first purified on a Ni–nitrilotriacetic acid (NTA) column (QIAGEN, Valencia, CA). The eluate was further purified on glutathione–Sepharose 4B resin (GE Healthcare, Piscataway, NJ). After TEV cleavage at 4°C overnight, the sample was concentrated and then loaded onto a HiLoad 16/60 Superdex 200 column (GE Healthcare) in 20 mM Tris at pH 8.0, 200 mM NaCl, and 0.3 mM tris(2- carboxyethyl)phosphine (TCEP). The sample fractions were pooled together, adjusted to 30 mM imidazole, and passed through 1 mL of glutathione–Sepharose 4B and then onto a Ni-NTA column (QIAGEN) to capture the residual GST- and His-tag fragments. The sample was adjusted to 20 mM Tris at pH 8.0, 200 mM NaCl, and 0.3 mM TCEP by buffer exchange in the concentrator, supplemented to 10% glycerol and flash frozen for long-term storage.

Myristoylated Nef constructs were co-expressed with human NMT in BL21 (DE3) Star cells and induced with 0.3 mM IPTG at 28°C for 8 hours. The media was supplemented with 50uM myristic acid, solubilized in ethanol, 30min prior to IPTG induction. The cell pellet was lysed by sonication and purified on a Ni-NTA column in 50 mM Tris at pH 8.0, 300 mM NaCl, 20 mM imidazole, 3 mM β-ME, and 0.5 mM phenylmethanesulfonyl fluoride (PMSF). Proteins were eluted with 300 mM imidazole and further purified by ion exchange (HiTrap Q HP). Enriched fractions were pooled and loaded onto a HiLoad 16/60 Superdex 75 column (GE Healthcare) in 20 mM Tris at pH 8.0, 200 mM NaCl, 5 mM MgCl2 and 0.3 mM TCEP. High-purity protein fractions were pooled and proteins were quantified by molar absorption measurements. Myristoylation was confirmed by whole-protein mass-spectrometry.

Myristoylated Arf1 constructs were co-expressed with yNMT in BL21 (DE3) Star cells and induced with 0.3 mM IPTG at 25°C overnight. The media was supplemented with 50uM myristic acid, solubilized in ethanol, 30min prior to IPTG induction. The cell pellet was lysed by sonication, clarified by centrifugation then filtered to 0.45um and passed over Q HP Sepharose resin. Myristoylated Arf1 was enriched by ammonium sulfate precipitation (35% saturation) then applied to HiLoad 16/60 Superdex 75 column (GE Healthcare) in 20 mM Tris at pH 8.0, 200 mM NaCl, 5 mM MgCl2 and 0.3 mM TCEP. Myristoylated and non-lipidated Arf1 proteins were separated by ion exchange on a monoS 10/100 GL with a linear salt gradient in MES pH 6.0 buffer supplemented with 1mM MgCl_2_ and 1mM DTT. Proteins were buffer exchanged into 20mM Tris pH 7.5, 100mM NaCl, 1mM MgCl_2_,1mM DTT and 10% glycerol with spin concentrators prior to storage at -80C. Proteins were quantified by molar absorption measurements. Myristoylation was confirmed by whole-protein mass-spectrometry.

MHC-I peptide was expressed in BL21 (DE3) Star cells and induced with 0.3 mM IPTG at 20°C overnight. The cell pellet was lysed by sonication and purified on a Ni-NTA column in 50 mM Tris at pH 8.0, 300 mM NaCl, 20 mM imidazole, 3 mM β-ME, and 0.5 mM phenylmethanesulfonyl fluoride (PMSF). The eluate was further purified on glutathione–Sepharose 4B resin (GE Healthcare, Piscataway, NJ). After TEV cleavage at 4°C overnight, the sample was recaptured on Ni-NTA resin, eluted, then loaded onto a HiLoad 16/60 Superdex 750 column (GE Healthcare) in 10 mM Tris at pH 7.0, 10 mM NaCl, 0.1mM EDTA and 1mM β-ME. Pure fractions were concentrated by evaporation on the SpeedVac then pooled and stored at -20°C. Clathrin triskelia was purified from food-grade porcine brain exactly as described (Jackson, 1993).

### Lipopeptide coupling

Peptides were treated with DTT for 1 hour at 25°C prior to buffer exchange on a PD-10 column (cytiva) according to the manufacturer’s instructions. Peptide concentration was quantified by molar absorption measurements. Peptide fractions were pooled then incubated at a molar ratio of 1:1.1 (excess lipid) with MPB-PE (Avanti Polar Lipids) for 3- 12 hours at 25C with end-over-end mixing. MPB-PE was vacuum desiccated in an amber glass vial prior to coupling. Lipopeptides were purified from unreacted peptides by reverse-phase liquid chromatography using C18 columns as previously described (Nickel and Wieland, 2001). Lipopeptide-containing fractions were dried under vacuum then resuspended in 2:1 chloroform/methanol and stored at -80°C. Lipopeptide concentrations were estimated by Coomassie stain on SDS-PAGE gels using peptide standards.

### Liposome preparation

Lipids were mixed at a ratio of 52% DOPC, 26% POPE, 7% POPS, 2% PI(4)P, 10% cholesterol and 3% lipopeptide for liposome experiments. Liposomes were prepared by resuspending the lipid mix to 1mg/ml lipid concentrations in HKM buffer (20mM HEPES pH 7.4, 125mM potassium acetate, 1mM DTT, 2mM MgCl_2_), freeze-thawed >5 times in liquid nitrogen and 42°C water bath with intermittent vortexing then extruded through a 0.05-μm polycarbonate filter (Avanti Polar Lipids) 11 times.

### Cryo-electron microscopy/tomography sample preparation

For AP-1^core^ cryoEM/ET samples, 2μM AP-1^core^, 8μM Arf1_myr_, 8μM Nef_myr_, 125nM ARNO, 200μM GTP were incubated with 0.25 mg/ml extruded liposomes for 1 hour at 25°C in HKM buffer prior to vitrification. For AP-1^βFL^ cryoEM/ET samples, 0.5μM AP-1^core^, 2μM Arf1_myr_, 2μM Nef_myr_, 125nM ARNO, 200μM GTP and 250nM clathrin were incubated with 0.125 mg/ml extruded liposomes for 1 hour at 25°C in HKM buffer prior to vitrification. 10nm BSA-gold fiducial marker was buffer exchanged into HKM buffer and concentrated to 10X and was added 1:10 to the sample just prior to sample application on the grid. 3.5 microliters of the reaction plus fiducial was applied to glow-discharged lacey carbon grids (200 mesh, thick; Ted Pella) and incubated at 100% relative humidity and 25°C for one minute prior to double-sided blotting for 4-5 seconds and plunge freezing into liquid ethane using a Vitrobot.

### Tilt series data acquisition

Bidirectional tilt series (3° increment, +/- 60° tilt range) were collected from vitrified samples containing AP-1^core^, Arf1_myr_, Nef_myr_ and liposomes incorporated with MHC-I^cyto^- lipopeptide. Data was collected on an FEI Titan Krios electron microscope operated at 300 kV using a Gatan Quantum energy filter with a slit width of 25 eV and a K3 direct detector operated in CDS mode. The total exposure of ∼120 e^-^/Å^2^ was equally distributed between 61 tilts. 3-4 frame movies were acquired for each tilt. The details of data collection are given in Table S1. The selection of acquisition areas was guided by suitability for high-resolution tomographic data collection (i.e., vitreous ice quality, lack of crystalline ice contaminations, and intactness of the carbon support).

Dose-symmetrical tilt series acquisition (Dodonova et al., 2017) were collected on cryo-preserved grids containing AP-1^βFL^, Arf1_myr_, Nef_myr_, clathrin and liposomes incorporated with MHC-I^cyto^-lipopeptide. Data was collected on an FEI Titan Krios electron microscope operated at 300 kV using a Gatan Quantum energy filter with a slit width of 25 eV and a K3 direct detector operated in CDS mode. The total exposure of ∼120 e^-^/Å^2^ was equally distributed between 41 tilts. 3-4 frame movies were acquired for each tilt. The details of data collection are given in Table S1. The selection of acquisition areas was guided by suitability for high-resolution tomographic data collection (i.e., vitreous ice quality, lack of crystalline ice contaminations, and intactness of the carbon support) and obvious presence of tubulated membranes.

### Tomogram reconstruction

Image preprocessing and tomogram reconstruction were performed essentially as described in (Hutchings et al., 2018). The IMOD v. 4.10.3 package (Mastronarde and Held, 2017) was used to align frames in raw movies and correct for detector gain. Tilt series were excluded from further analysis based on large tracking errors or significant beam-induced sample movements. Defective high-tilt images (due to tracking error, large objects like a grid bar, or contaminations coming in the field of view) were also removed before tilt series alignment. Tilt series were aligned using gold fiducial markers in the IMOD package. CTFPLOTTER (within IMOD) was used to estimate the defocus. Aligned, unbinned tilt series were low-pass filtered according to cumulative dose (Grant and Grigorieff, 2015), 3D CTF corrected by phase-flipping using 15nm strip widths and reconstructed into tomograms by back-projection using NovaCTF (Turonova et al., 2017). Reconstructed tomograms were binned by factors of 2 and 4 using antialiasing in IMOD. Bin4 tomograms were SIRT-like filtered and used for tube annotation in Dynamo (Castano-Diez et al., 2012). Tomograms were denoised with IsoNet for illustration purposes only (Liu et al., 2021).

### Subtomogram averaging

Subtomogram alignment and averaging were done in Dynamo (1.1.532). MATLAB scripts were adapted and/or written in-house. Centers of coated tubules were manually traced in bin4, filtered tomograms, and their diameters were recorded in Dynamo (1.1.532). All tubules were annotated regardless of apparent diameter. The positions of subtomograms were defined on the surface of tubes with uniform radial sampling and uniform separation between rings. Initial subtomogram orientations were set to be normal to the membrane surface. The full dataset of annotated tubes was partitioned into 10nm windows (10- 20nm, 20-30nm, 30-40nm, etc.) and each subset was individually processed to identify correlations in AP-1 lattice structure and tube diameter.

For each subset, two thousand random subtomograms were extracted from bin4, unfiltered tomograms and averaged according to their initial orientations, producing an initial reference. The subtomograms were then aligned to this reference, allowing both shifts and angular search. A saddle-shaped mask was applied passing the protein layer and the membrane. Eight iterations of such alignment were performed with a 20° to 2° angular search increment and an appropriate low-pass filter. A second round of alignments were performed using two, noisy versions of the resulting average plus low-pass filtering to parse γ - γ lattice-centered subtomograms and β1-β1 lattice-centered subtomograms. The resulting two averages were shifted and rotated to more precisely center the γ - γ or β1-β1 tracks before further processing.

The resulting averages (γ - γ centered lattice or β1-β1 centered lattice) were used as templates to align the complete 10nm-window datasets, using alignment and sampling parameters as above. Overlapping subtomograms resulting from oversampling at the initial extraction stage were removed by selecting the subtomogram with the highest cross-correlation score within a distance threshold equivalent to a third of the box size. Subtomograms were then split into odd and even half datasets for further processing. Subsequent alignments were performed independently on the odd and even halfsets. The search space and angular increments were gradually decreased, and the low-pass filter was gradually moved toward higher resolution. After each round of alignments, subtomogram cross-correlation values were reweighted as a function of the θ Euler angle, as in (Hutchings et al., 2018), on a tube-by-tube basis. Cross-correlation cutoff values were determined from visual inspection of the subtomogram real-space coordinates colored by cross-correlation values using the chimera PlaceObject plugin . Thresholds were applied globally to both half-sets. Subvolumes located close to the edge of tomograms were also removed. At the end of each iteration, the resolution of a given structure was estimated by Fourier shell correlation of the independent half-maps in Dynamo.

Subboxing was performed in Dynamo (1.1.532). Coordinates for individual dimers or specific interfaces mapped to positions on the γ - γ centered lattice or β1-β1 centered lattice and used extract to subtomograms, initially from bin4 tomograms. Subtomogram alignment and averaging of each subboxed region was performed as described in ‘Subtomogram Alignment’. The global resolution of each EM map was estimated in Relion-3.1 (Zivanov et al., 2018). Local resolution was measured using relion_postprocess from the Relion 3.1 package (Zivanov et al., 2018). The local resolution ranges for each map are listed in Table S2. The reported global resolution is based on FSC 0.143 values. Final half-maps were cropped to remove density corresponding to areas residing outside the mask used for subtomogram alignment. The final sharpened maps were prepared using local resolution filtering and denoising implemented in Relion 3.1.

### Model building

Rigid-body fitting and molecular visualization were performed in Chimera (1.14) and ChimeraX (1.3) (Pettersen et al., 2021). To build models of the asymmetric unit on narrow tubes, a single protomer from the trimeric AP-1:Arf1:BST2-Nef [Protein Data Bank (PDB) 6CM9] assembly was fit to the STA map. To model the MHC-I^cyto^ peptide bound to Nef, the crystal structure of μ1:Nef-MHC-I (PDB:4EN2) was fit to the same STA map and a composite atomic model was created by replacing the MHC-I peptide and Nef of 4EN2 with the corresponding models in 6CM9. To model the N-terminus of Arf1 (2-16), the full-length, human Arf1 model produced by AlphaFold2 (Uniprot: P84077) was aligned to Arf1^γ^ in the new composite model and the N-terminal residues were manually adjusted to fit the STA map density in ISOLDE. The modeled N-terminus was then cropped and stitched to the Arf1^γ^ from 6CM9 in Coot (version 0.9.1). The full-length modeled Arf1 replaced Arf1^β1^ and Arf1^γ^ models in the new composite structure. To build the asymmetric unit on wide tubes, the asymmetric unit from narrow tubes was fit to the STA map corresponding to the asymmetric unit on wide tubes. One additional Arf1 molecule from the narrow asymmetric unit was manually placed in the density associated with Arf1^γ’^. Composite maps were generated for visualization purposes only. Overlapping regions of the γ−γ and β1-β1 lattices were overlayed and combined in Chimera (1.14) or ChimeraX (1.3) using the vop command. Helical parameters for the lattice were calculated by hand using the composite maps. Models were real space refined against merged, locally filtered maps by global minimization in Phenix (1.20-4459) and global FSC 0.143 as the resolution cut- off (Liebschner et al., 2019).

### Modeling the open AP-1:Arf1 lattice

Previously, a model of an open AP-1:Arf1 hexagonal lattice compatible with a hexagonal clathrin coat was generated by combining open AP-1:Arf trimers (Shen et al., 2015) and crystal structure of an Arf1^β^-linked dimer (Ren et al., 2013). We revisited this model on the basis of the membrane-associated Arf1 dimers visualized in this study. Assembly of AP-1:Arf1 trimers linked by ‘narrow’ tube Arf1^γ^:γ dimeric interfaces results in a pentameric arrangement of AP-1 molecules that matches the spatial distribution of clathrin hubs at pentagonal faces of clathrin cages (Fig. S5). Assembly with Arf1^γ^:γ dimeric contacts like those in the ‘wide’ tubular lattice results in a flatter and wider arrangement of AP-1 molecules such that a hexagon results instead (Fig. S5). The binding site for the third Arf1 molecule remains intact as observed in the wide tube lattice and the dimensions of the hexagon again match the dimensions of the clathrin hexagonal face (Fig. S5).

### Sequence alignments

Primary sequences encoding Arf and AP proteins were aligned using the COBALT, clustal omega and/or Tcoffee/Mcoffee/Espresso Server. Alignments were formatted in boxshade.

### DNA plasmids and siRNA for cell imaging

The pcDNA5 NA7-Nef-HALO vector was generated by fusing the codon optimized NA7- Nef sequence (protein ID ABB51086.1), SGGTG linker – HALO tag DNA sequence, and the pcDNA5 vector backbone via Gibson assembly. The pCI NL4-3-Nef-HALO vector was generated by fusing the pCI NL4-3 Nef and STSGGTG linker - HALO tag DNA sequence via Gibson assembly. pEGFPN1 AP1-GFP was generated by inserting the Hs AP1M1 sequence with GSGS linker into a pEGFPN1 backbone through double restriction digest with XhoI and HindIII. Clathrin knockdown was conducted with SMARTPool ON-TARGETplus Human CLTC siRNA purchased from Dharmacon.

### Tissue culture

All live-cell imaging was conducted using HeLa cells obtained from the UCB Cell Culture Facility. Cells were cultured in DMEM F12 media supplemented with 10% FBS and PenStrep, and grown at 37°C and 5% CO_2_. 24 hours prior to live-cell imaging, cells were plated onto 8 well chambered cover glass (Cellvis C8-1.5H-N) and incubated with 100nM JF635 dye. Before imaging, cells were transferred to imaging media consisting of DMEM F12 -phenol red supplemented with 10% FBS and PenStrep. Cells were allowed to recover for at least 10 minutes in the incubator prior to movie acquisition.

### DNA Transfection and siRNA Knockdown

For knockdowns, HeLa cells were cultured to ∼60% confluency in 6-well plates and transfected with Lipofectamine 3000 (Invitrogen L3000) following directions provided by the manufacturer. The final concentration of siRNA used for knockdown was 20nM. After one day, cells were split into 12-well plates for transfection with DNA plasmids. The following day, 1ug of each DNA plasmid (AP1-GFP, NA7-Nef-HALO, and/or NL4-3-Nef- HALO) was transfected as desired using the *TransIT-*LT1 reagent (MIR2304) following directions provided by the manufacturer. Finally, cells were plated onto the 8 well chambered cover glass for imaging.

### Cell lysis and Western blots

HeLa cells, grown in 12-well plates, were washed with PBS, trypsinized, and resuspended in 24ul of lysis buffer, consisting of 50mM Hepes pH 7.4, 150mM NaCl, 1 mM MgCl_2_, 1% NP40, PhosSTOP phosphatase inhibitor (Roche 04906845001), and protease inhibitor (Roche 1183617001). Cells were lysed for 15 minutes on ice with gentle agitation every 5 minutes. Lysed cells were then centrifuged at 4°C and 13000 rpm for 15 minutes and 18ul of the supernatant was collected as lysate. The lysate was mixed with Laemmeli reducing buffer with 5% β-mercaptoethanol and 10ul of the final solution was loaded onto a 10% acrylamide gel for SDS-PAGE analysis. The SDS-PAGE gel was transferred onto nitrocellulose membrane for immunoblotting via wet transfer. Blots were blocked with 5% milk in TBS and incubated with primary antibody. The antibodies used for western blotting was anti-cltc (1:1000 in 0.1% Tween20, 0.05% milk in PBS, RT 1h, Abcam ab21679) and anti-GAPDH (1:10000 in TBST, RT 1 hr, Abcam ab9485). Blots were then incubated with IRDye 800/680 conjugated antibodies (1:10000 in 5% milk TBST, RT 1hr, 926-32212 926-68071) and imaged on the LICOR Odyssey scanner.

### Cell imaging

Live-cell imaging was conducted on a Zeiss LSM900 with Airyscan 2.0 detection in SR- 2Y mode, using the Zen Blue software. All imaging was conducted in an incubation chamber at 37°C and 5% CO_2_. 10um Z-stack movies with 140nm intervals were acquired for all analyzed cells. During multi-channel imaging, all channels were frame-sequentially acquired before moving to the next Z-slice to minimize crosstalk and time offset. Images were batch processed by ZEN Blue software for Airyscan reconstruction using filter strength setting 5 and 3D processing.

### Image analysis

To test for colocalization between Nef and AP1 signal, both channels from the same slice in a Z-stack was analyzed. Both channels were first background corrected using the 15 pixel median filtering as described above, and thresholded using the “Auto Threshold” Otsu and Li thresholds for the AP1 and Nef channels respectively in imageJ. Particles less than 0.10 μm^2^ were filtered out. AP1 signal was then classified as “spheroid” if particles had circularity between 0.75 – 1.00 or as “tubule” if particles had circularity between 0 – 0.75 according to the “Analyze Particles” function. Using the thresholded images of AP1 and Nef, the Manders correlation coefficient was calculated for each type of AP1 structure using the JACoP plugin in imageJ (Bolte and Cordeliès 2006). Specifically, the fraction of AP1 pixels overlapping with Nef pixels was reported for colocalization. To confirm that colocalization of AP1 and Nef was not random, the Van Steensel CCF function was generated using JACoP as well. The imageJ scripts used for AP1 tubule analysis can be found here: https://github.com/yuichiro-iwamoto/AP1_tubule_quantification

### Statistical analysis

Statistical analysis of AP1/Nef colocalization was performed using the GraphPad Prism 9 software. In all cases, the Kruskal-Wallis test was used to make one-way comparisons across all conditions. ns: p > 0.05, *: p ≤ 0.05, **: p ≤ 0.01, ***: p ≤ 0.001, ****: p ≤ 0.0001.

---

**Supplementary Figure 1.**
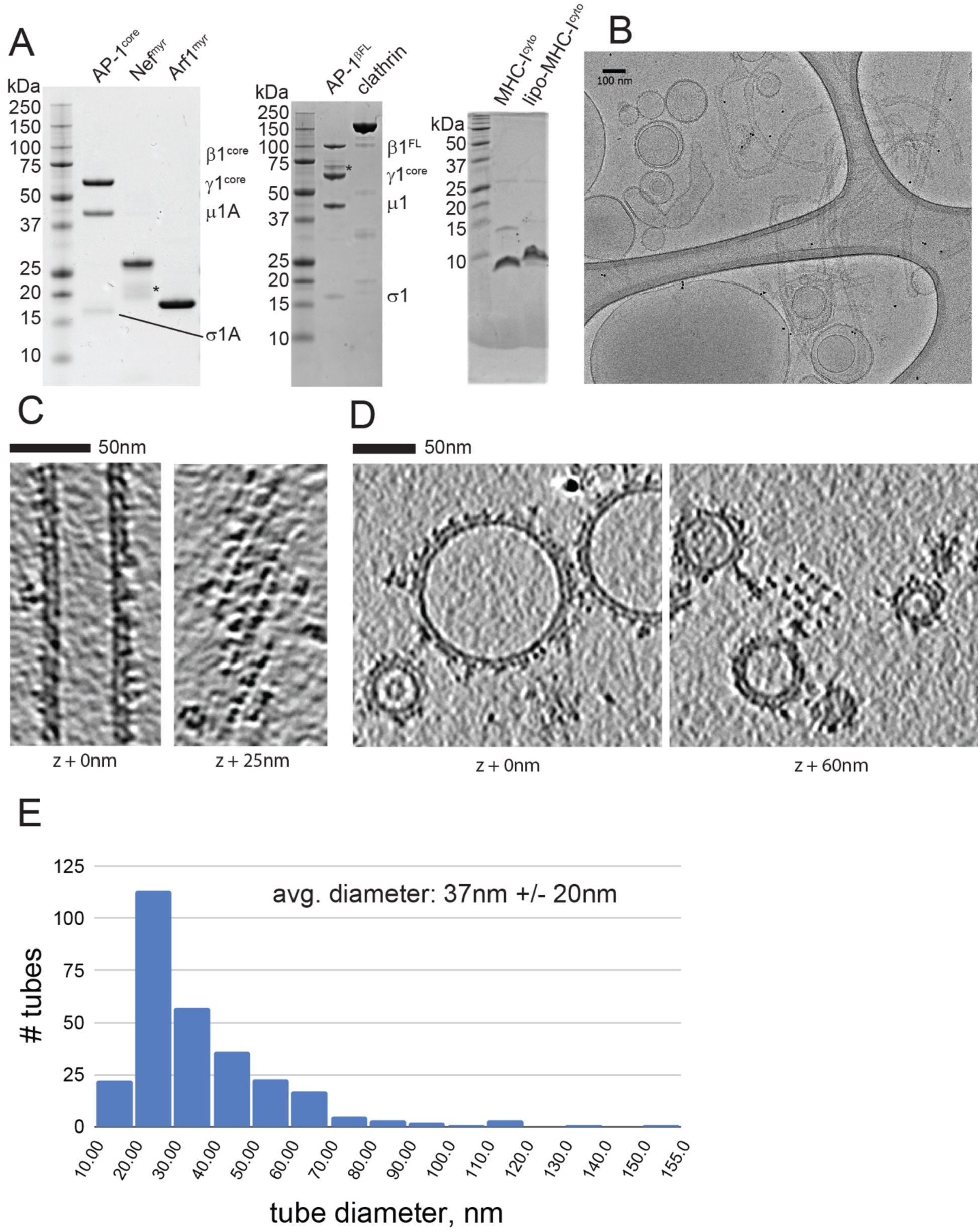
(A) SDS-PAGE gel stained with Coomassie showing purity of proteins and lipopeptide reagents used in the study. Asterisks denote degradation products. (B) Sample cryoEM image of AP-1^bFL^, Arf1_myr_, Nef_myr_ mixed with extruded liposomes incorporated with MHC-I^cyto^-lipopeptide and visualized on lacey holey carbon grids. (C) Representative cryo-electron tomographic slices through an AP-1^core^:Arf1_myr_:Nef_myr_ coated tube incorporated with MHC-I^cyto^ lipopeptide. (D) Representative cryo-electron tomographic slices through an AP-1^core^:Arf1_myr_:Nef_myr_ coated vesicle incorporated with MHC-I^cyto^ lipopeptide. (E) Distribution of tube diameters from the dataset used for subtomogram averaging and structure determination. 284 tubes were annotated from 72 tomograms containing AP-1^bFL^, Arf1_myr_, Nef_myr_ coated membranes incorporated with MHC-I^cyto^-lipopeptide.

**Supplementary Figure 2.**
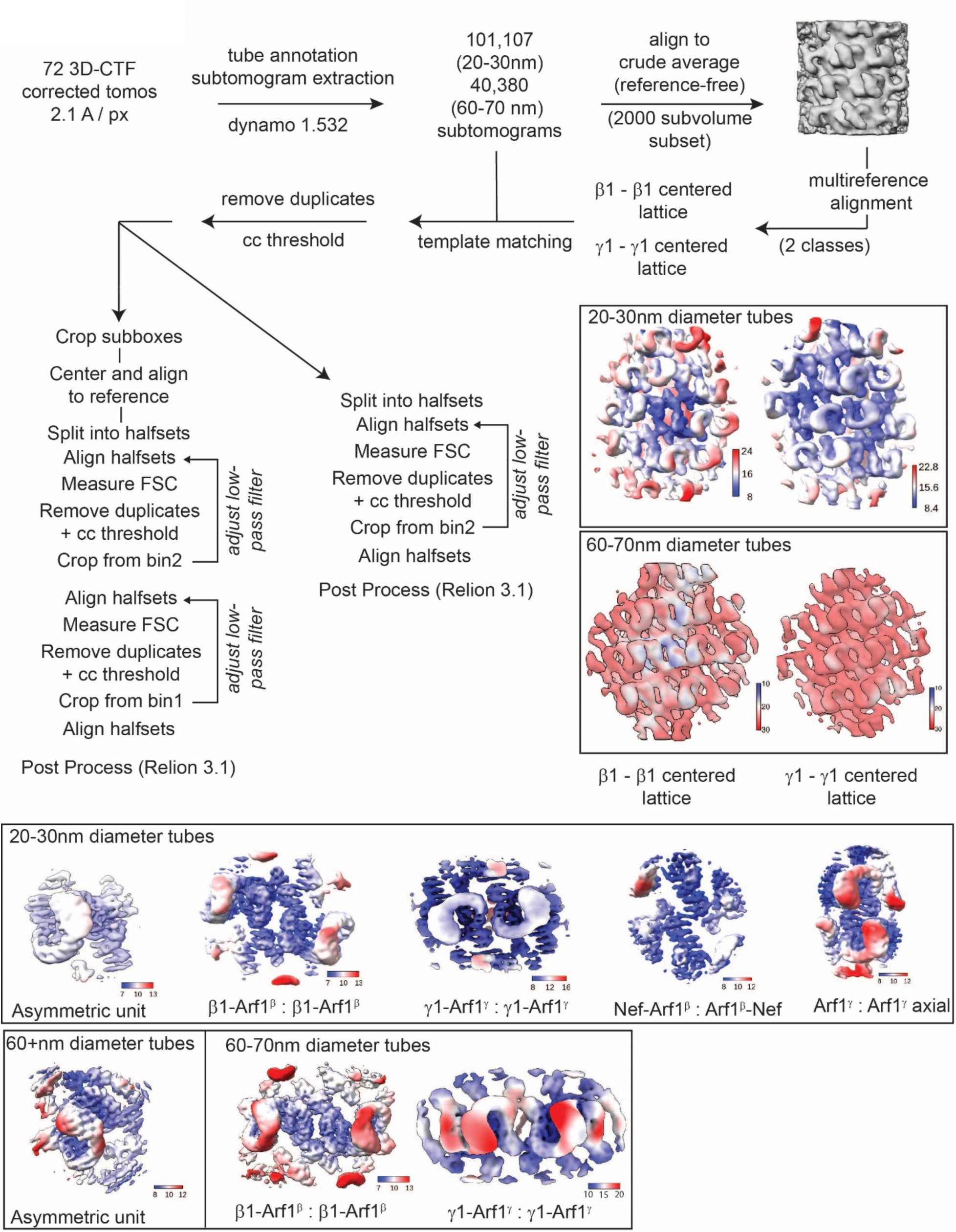
Subtomogram averaging workflow.

**Supplementary Figure 3.**
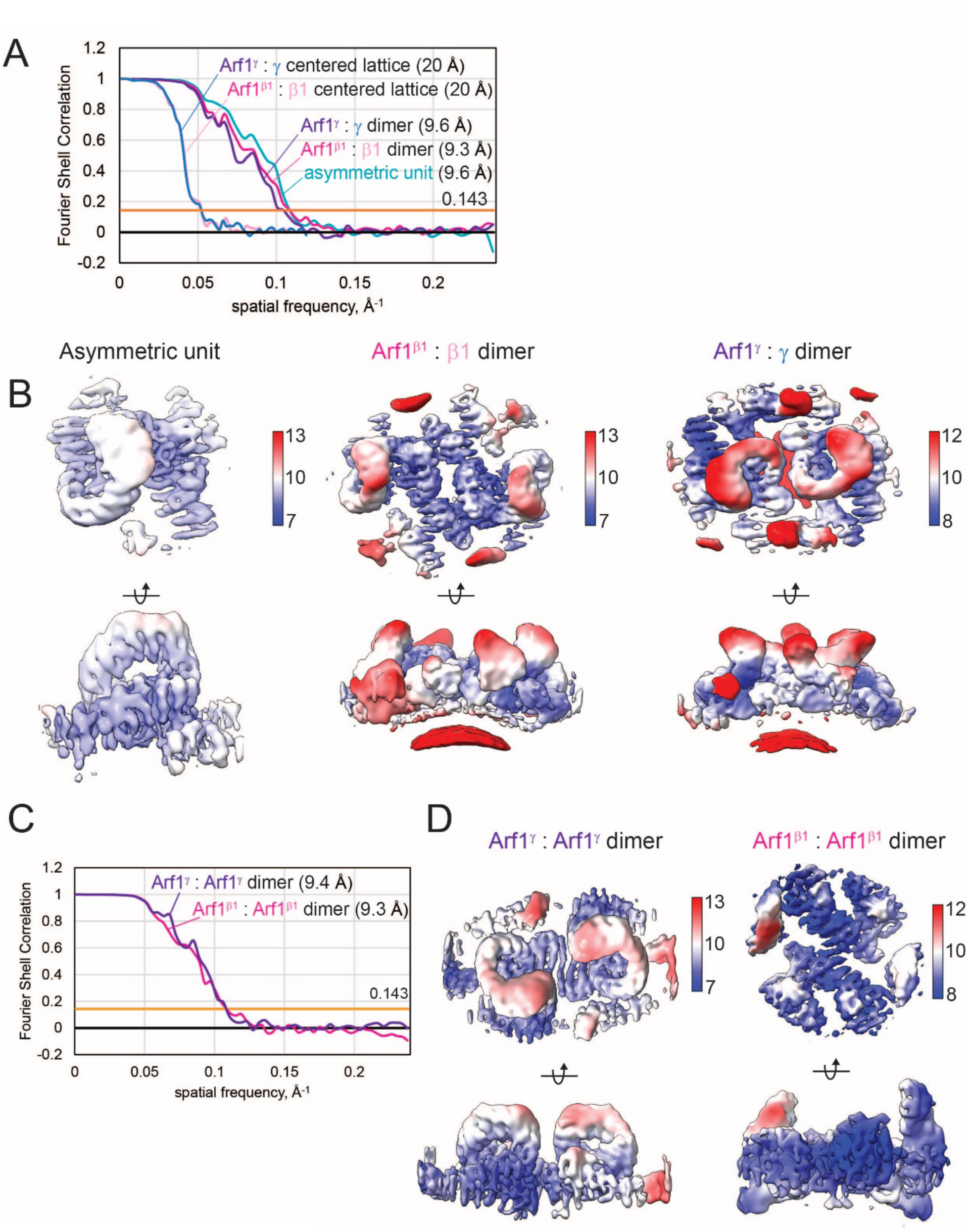
Global and local resolution of individual EM maps derived from subtomogram averaging (A) FSC curves for maps shown in (B). (B) Local resolution-filtered maps of structures determined on narrow tubes. (C) FSC curves for maps shown in (D). (B) Local resolution-filtered maps of structures determined on narrow tubes, continued.

**Supplementary Figure 4.**
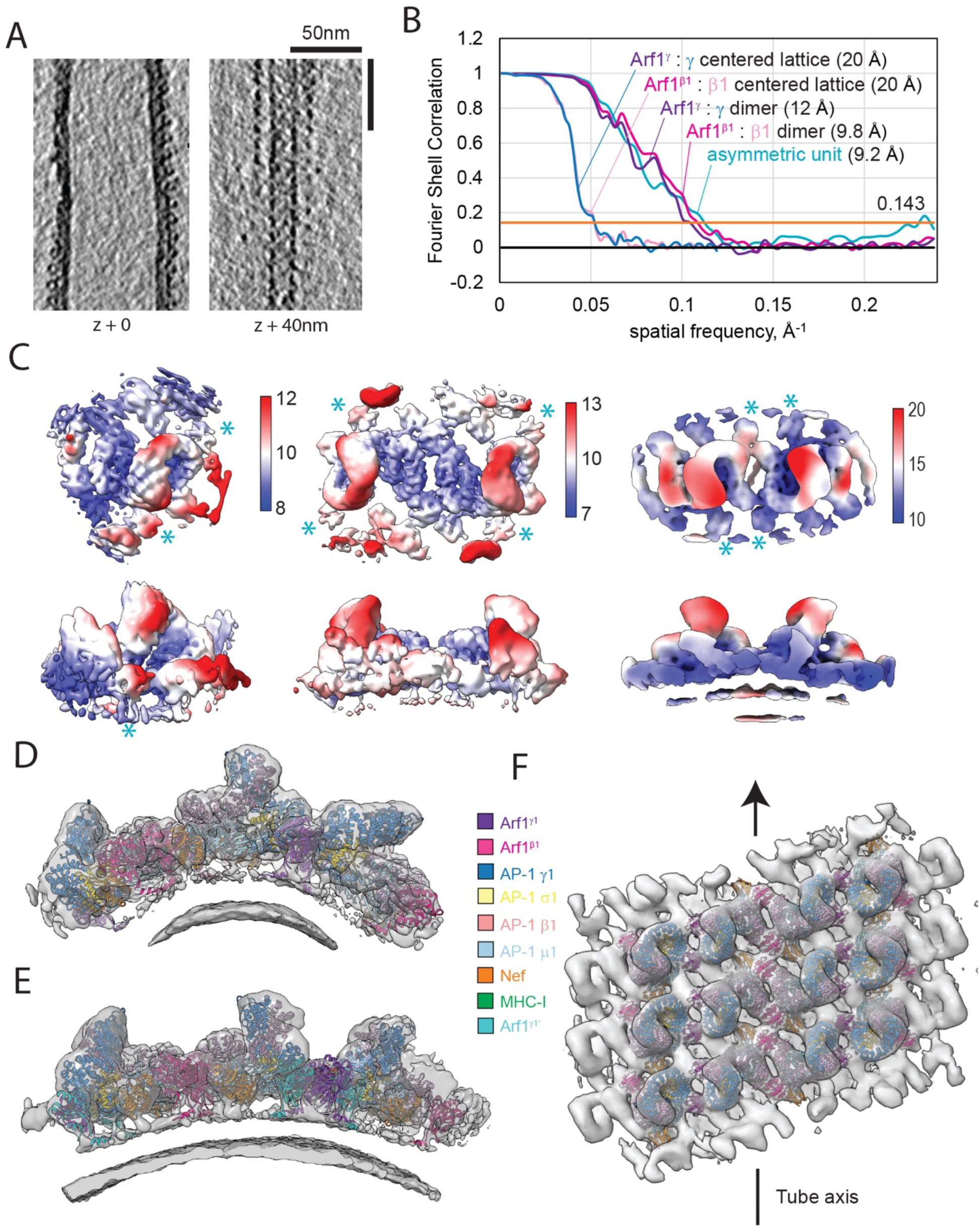
(A) Representative cryo-electron tomographic slices through an AP-1^bFL^:Arf1_myr_:Nef_myr_ coated wide tube (>60nm diameter) incorporated with MHC-I^cyto^ lipopeptide. (B) Global FSCs of individual EM maps derived from subtomogram averaging of particles on wide tubes. (C) Local resolution estimates of individual EM maps derived from subtomogram averaging of particles on wide tubes. (D) and (E) Composite maps and corresponding atomic models of three AP-1 / Arf1 / Nef asymmetric units laterally engaged by Arf1^β1^ : β1 and Arf1^γ^ : γ dimeric interfaces. Maps are shown as side-on views. (D) map and model on narrow tubes. (E) map and model on wide tubes. (F) Composite map of the γ : γ -centered lattice and β1 : β1-centered lattice on wide tubes, shown as top view

**Supplementary Figure 5.**
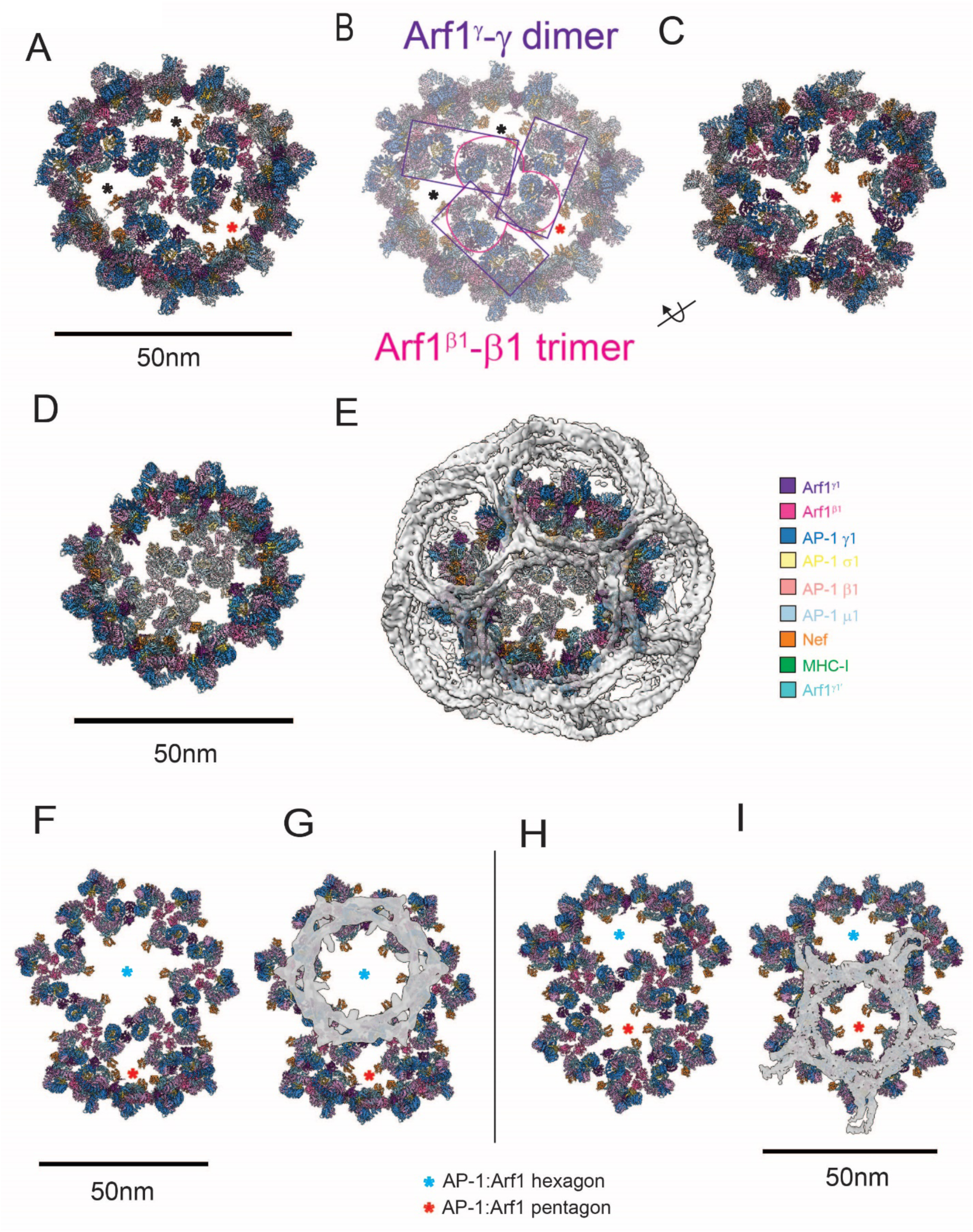
Model of an AP-1:Arf1 hexagonal and pentagonal lattice (A) A hypothetical clathrin coat-like assembly of AP-1:Arf1:Nef complexes. Arf1^β1^-linked AP-1 trimers are bridged by Arf1^γ^ - AP-1 γ dimeric contacts observed in the AP- 1:Arf1:Nef lattice on narrow tubes. Asterisks denote pentagonal faces. (B) Outlines of the Arf1^β1^-linked AP-1 trimers (pink) and Arf1^γ^ : g -linked AP-1 dimers (purple) overlaid on the model shown in (A). Asterisks denote pentagonal faces. (C) Rotated view of the model shown in (A). The pentagonal face denoted by a red asterisk is the same as shown in (A). Nef is bound to μ1 domains along the inner face of the pentagon resulting in the projection of its dileucine loop towards the center of the hole. (D) A hypothetical clathrin coat-like assembly of AP-1:Arf1:Nef complexes using a mix of narrow and wide Arf1^γ^ - AP-1 γ dimeric contacts. (E) The structure shown in (D) overlaid with a complete clathrin cage (EMDB: 0118) (F) A hypothetical clathrin coat-like assembly of AP-1:Arf1:Nef complexes. Arf1^β1^-linked AP-1 trimers bridged by Arf1^γ^ - AP-1 γ dimeric contacts observed in the AP-1:Arf1:Nef lattice on wide tubes result in a hexagonal arrangement (blue asterisks). Arf1^β1^-linked AP-1 trimers bridged by Arf1^γ^ - AP-1 γ dimeric contacts observed in the AP-1:Arf1:Nef lattice on narrow tubes result in a pentagonal arrangement (red asterisks). (G) The structure shown in (F) overlaid with a hexagonal face of a clathrin cage (EMDB: 12980) (H) Rotated view of (F) such that the pentameric face is centered. (I) The structure shown in (H) overlaid with a pentagonal face of a clathrin cage (EMDB: 21615)

**Table S1.** Cryo-electron tomography data collection

|  | AP-1<sup>core</sup>, Arf1<sub>myr</sub>, Nef<sub>myr</sub><br>+ MHC-I<sup>cyto</sup>-liposomes | AP-1<sup>βFL</sup>, Arf1<sub>myr</sub>, Nef<sub>myr</sub>, clathrin<br>+ MHC-I<sup>cyto</sup>-liposomes |
| --- | --- | --- |
| Grids | Lacey Holey Carbon (200 mesh, Thick carbon – Ted Pella) | Lacey Holey Carbon (200 mesh, Thick carbon – Ted Pella) |
| Cryo-specimen freezing | Vitrobot Mark IV | Vitrobot Mark IV |
| Microscope, Voltage (keV) | Titan Krios, 300 | Titan Krios, 300 |
| Detector | Gatan Quantum K3 | Gatan Quantum K3 |
| Energy filter slit width (eV) | 25 | 25 |
| Electron exposure (e<sup>-</sup> / Å<sup>2</sup>) dose fractionation | ~120, uniformly distributed over tilt series | ~120, uniformly distributed over tilt series |
| Defocus range (μm) | -6 μm | -1.5 μm to -4.5 μm |
| Tilt scheme (min/max, step) | -60°/+60°, 2°, bidirectional | -60°/+60°, 3°, dose-symmetrical (Hagen scheme) |
| Movie recording | 5-6 frames per tilt | 4-5 frames per tilt |
| Magnification (times) | X26,000 | X42,000 |
| Pixel size | 3.07 Å / pixel | 1.05 Å / pixel (super resolution) |
| Tomograms acquired | 4 | 72 |
| Traced membranes (no.) and luminal diameter (mean +/- std) |  | 284 tubes, 37 nm +/- 20 nm average luminal diameter |

**Table S2.** Subtomogram averaging, narrow tubes (20-30nm diameter)

| | $\beta 1$-<br>Arf1: $\beta 1$:Arf1<br>centered<br>lattice | $\gamma 1$-<br>Arf1: $\gamma 1$:Arf1<br>centered<br>lattice | $\gamma 1$-<br>Arf1: $\gamma 1$:Arf1<br>lateral dimer | $\beta 1$-<br>Arf1: $\beta 1$:Arf1<br>lateral dimer | $\gamma 1$-<br>Arf1: $\gamma 1$:Arf1<br>axial dimer | Nef : Arf1<sup>$\beta 1$</sup><br>axial<br>interface | Asymmetric<br>unit AP-<br>1<sub>1</sub>:Arf1<sub>2</sub>:Nef |
| --- | --- | --- | --- | --- | --- | --- | --- |
| EMDB |  |  |  |  |  |  |  |
| PDB ID | n.a. | n.a. |  |  |  |  |  |
| Box Size / bin / pixel<br>spacing at bin | 128 / bin2 /<br>4.2 | 128 / bin2 /<br>4.2 | 128 / bin1 /<br>2.1 | 128 / bin1 /<br>2.1 | 128 / bin1 /<br>2.1 | 128 / bin1 /<br>2.1 | 96 / bin1 /<br>2.1 |
| STA alignments and<br>averaging | Dynamo | Dynamo | Dynamo | Dynamo | Dynamo | Dynamo | Dynamo |
| Initial subtomograms<br>(no.) / after duplicate<br>removal | 27877 | 23154 | 29941 | 20560 | 37976 | 43110 | 61864 |
| Final subtomograms<br>(no.) | 6141 | 8144 | 7343 | 7004 | 8615 | 8174 | 16422 |
| Tomograms (initial /<br>final) | 38 / 35 | 38 / 38 | 38 / 38 | 38 / 36 | 38 / 38 | 38 / 38 | 38 / 31 |
| Tubes (initial / final) | 131 / 96 | 131 / 129 | 131 / 119 | 131 / 96 | 131 / 125 | 131 / 124 | 131 / 70 |
| Symmetry imposed | C1 | C1 | C2 | C2 | C2 | C2 | C1 |
| Global resolution -<br>FSC threshold 0.143,<br>Å (Relion 3.1) | ~20 | ~20 | 9.6 | 9.3 | 9.4 | 9.3 | 9.6 |
| Resolution range (Å) |  |  | 8-12 | 7-13 | 7-13 | 7-13 | 7-13 |

**Table S3.** Subtomogram averaging, wide tubes (60-70nm diameter or >60nm diameter***)

| | $\beta 1$-Arf1: $\beta 1$:Arf1<br>centered lattice | $\gamma 1$-Arf1: $\gamma 1$:Arf1<br>centered lattice | $\gamma 1$-Arf1: $\gamma 1$:Arf1<br>lateral dimer | $\beta 1$-Arf1: $\beta 1$:Arf1<br>lateral dimer | Asymmetric unit<br>AP-<br>1<sub>1</sub>:Arf1<sub>3</sub>:Nef<sup>***</sup> | $\gamma 1$-<br>Arf1: $\gamma 1$:Arf1<br>lateral<br>dimer<sup>***</sup> |
| --- | --- | --- | --- | --- | --- | --- |
| EMDB |  |  |  |  |  | n.a. |
| PDB ID | n.a. | n.a. |  |  |  | n.a. |
| Box Size / bin / pixel<br>spacing at bin | 112 / bin2 / 4.2 | 112 / bin2 / 4.2 | 144 / bin1 / 2.1 | 144 / bin1 / 2.1 | 96 / bin1 / 2.1 | 48 / bin4 /<br>8.4 |
| STA alignments and<br>averaging | Dynamo | Dynamo | Dynamo | Dynamo | Dynamo | Dynamo |
| Initial subtomograms<br>(no.) / after duplicate<br>removal | 4208 | 4208 | 8601 | 6943 | 53025 / 25534 | 98569 / |
| Final subtomograms<br>(no.) | 2719 | 2454 | 5094 | 5219 | 18183 | 9066 |
| Tomograms (initial /<br>final) | 13 / 12 | 13 / 12 | 13 / 13 | 13 / 12 | 24 / 24 | 24 / 24 |
| Tubes (initial / final) | 17 / 16 | 17 / 16 | 17 / 17 | 17 / 16 | 33 / 31 | 33 / 33 |
| Symmetry imposed | C1 | C1 | C2 | C2 | C1 |  |
| Global resolution -<br>FSC threshold 0.143,<br>Å (Relion 3.1) | ~20 | ~20 | 11.6 | 9.8 | 9.2 | ND |
| Resolution range (Å) |  |  | 10-20 | 7-13 | 8-12 |  |

**Table S4.** Helical parameters of AP-1 lattice on narrow and wide tubes

|  | 20-30nm diameter | 60-70nm diameter |
| --- | --- | --- |
| rise | 0.95 nm (calc) | 3.25 nm (calc) |
| pitch | 10 nm | 70 nm |
| units per turn | 10.5 | 21.5 |
| twist | 34° | 16-17° |

